# Movement-related increases in subthalamic activity optimize locomotion

**DOI:** 10.1101/2023.12.07.570617

**Authors:** Joshua W. Callahan, Juan Carlos Morales, Jeremy F. Atherton, Dorothy Wang, Selena Kostic, Mark D. Bevan

**Affiliations:** Department of Neuroscience, Feinberg School of Medicine, Northwestern University, Chicago, IL 60611, USA

**Keywords:** basal ganglia, motor control, indirect pathway, hyperdirect pathway, action execution, gait, Huntington’s disease, Parkinson’s disease, neuromodulation, deep brain stimulation

## Abstract

The subthalamic nucleus (STN) is traditionally thought to restrict movement. Lesion or prolonged STN inhibition increases movement vigor and propensity, while optogenetic excitation typically has opposing effects. Subthalamic and motor activity are also inversely correlated in movement disorders. However, most STN neurons exhibit movement-related increases in firing. To address this paradox, STN activity was recorded and manipulated in head-fixed mice at rest and during self-initiated and -paced treadmill locomotion. The majority of STN neurons (type 1) exhibited locomotion-dependent increases in activity, with half encoding the locomotor cycle. A minority of neurons exhibited dips in activity or were uncorrelated with movement. Brief optogenetic inhibition of the dorsolateral STN (where type 1 neurons are concentrated) slowed and prematurely terminated locomotion. In Q175 Huntington’s disease mice abnormally brief, low-velocity locomotion was specifically associated with type 1 hypoactivity. Together these data argue that movement-related increases in STN activity contribute to optimal locomotor performance.

## Introduction

The glutamatergic subthalamic nucleus (STN) is a small but key component of the basal ganglia, a group of subcortical brain nuclei critical for habitual/automatic, goal-directed/flexible, and motivated behaviors^1^. The STN is classically thought to prevent, limit, or stop movement^2–9^. At rest, tonic driving of inhibitory basal ganglia output by the STN may suppress the activity of multiple motor centers^2^. In addition, hyperdirect and indirect pathway-mediated elevations in STN activity have been proposed to prevent an imminent action, terminate execution of an action, and/or facilitate execution of volitional movement through suppression of competing actions^3–14^. Consistent with these functions 1) lesions or prolonged pharmacological, optogenetic or chemogenetic inhibition of the STN leads to dyskinesia, hyperkinesia, premature or inappropriate responding, and stereotyped behaviors, such as excessive grooming^5,13,15–18^ 2) brief or prolonged optogenetic excitation of the STN typically prevents, reduces, or terminates movement^7,9,11,17,19^, but see^20,21^ 3) a subset of STN neurons exhibit elevated activity during passive or voluntary movement, whereas the activity of a partially overlapping subset of neurons is linked to stop, no go, or switch signaling^6–14,22–27^. Together, these data argue that the STN can suppress movement in some contexts but whether the STN actively facilitates the execution of volitional actions is less clear^13,28–30^.

The encoding properties of individual STN neurons are related to the cortico-basal ganglia thalamo-cortical loop(s) in which they reside^8–11,13,23,24,26,27,31^. Thus, movement-related activity is more prevalent in the dorsal and lateral STN, which is preferentially innervated by the motor cortex, and exhibits a somatotopic organization consistent with the anatomical arrangement of cortical afferents^10,13,14,23,24,26,31–41^. In contrast, no go, stop, limbic signaling is more common in the ventral and medial aspects of the STN, which are more strongly innervated by prefrontal, limbic, and higher order motor cortical areas^8,10,12,35,36,38,39,42^. However, some STN neurons exhibit complex combinations of encoding properties presumably due to partial overlap of functionally heterogeneous cortico-STN terminal fields and STN neuron dendrites that traverse distinct functional zones, especially in rodents^8,10,11,14,43^. Consistent with this general framework, lesions or pharmacological/chemogenetic inhibition of the motor subthalamic region increase movement or produce dyskinesia/ballism, whereas more medially placed manipulations generate stereotyped behavior^13,15–19,37^. In addition to their functional diversity, STN neurons exhibit heterogeneous molecular properties, e.g., parvalbumin (PV)- and calretinin-expressing STN neurons are concentrated in the motor and limbic zones of the nucleus, respectively, arguing that these neuron subtypes subserve distinct functions^44,45^. Furthermore, adjacent PV-expressing and non PV-expressing neurons in the dorsolateral STN have distinct connections and synaptic properties, arguing for additional STN functional subtypes and circuit complexity^46^.

Our understanding of STN function has been greatly informed by the impact of dysregulated STN activity in psychomotor disorders. In Parkinson’s disease (PD), STN hyperactivity and/or excessive synchronization have been linked to akinesia, rigidity, and gait deficits^2,27,47–49^. In contrast STN hypoactivity has been proposed to underlie ballism and to contribute to chorea, gait abnormalities, and psychiatric symptoms in Huntington’s disease^2,27,50–52^ (HD). Deep brain stimulation (DBS) of the dorsolateral STN ameliorates akinesia, rigidity, and gait deficits in PD^53,54^, whereas DBS of the ventromedial STN reduces obsessive compulsive behavior^55^ and can elicit affective and cognitive side effects in PD^56,57^, consistent with the STN’s regional, anatomically based functional organization.

Much of the work described above studied rodents and non-human primates during the execution of highly trained licking behaviors and reaching respectively, or human patients with neurologic/psychiatric disorders on and off therapy. The role of the STN in the execution of self-initiated, naturalistic behaviors like locomotion therefore remains an open question^28^. The effects of lesions or pharmacologic/optogenetic/chemogenetic manipulations to address a causative role for the STN in action execution or active stopping or the expression of disease symptoms have also been difficult to interpret due to their powerful, non-specific, prolonged, and irreversible or slowly reversible effects. Finally, given that the STN is a deep and highly vascularized structure, comprised of small, tightly packed neurons, well-isolated recordings of individual neurons are technically challenging^25^, especially under freely moving configurations. With these considerations in mind, we re-examined the role of the STN in motor control using silicon probe/optrode recordings in head-fixed mice at rest and during self-initiated, self-paced treadmill locomotion. Brief 5 second optogenetic inhibition was used to infer the impact of ongoing STN activity on rest or an already initiated locomotor bout. In addition, we compared STN encoding in wild type (WT) mice with Q175 HD mice because Q175 mice exhibit subtle but consistent gait deficits that are analogous to those in HD patients^58–61^ and may in part reflect the dysregulated activity of STN neurons in HD and its experimental models^50–52,62^.

## Results

1-2 weeks after surgically affixing a metal plate to the skull, WT and Q175 mice were habituated to head-fixation on a cylindrical or linear self-paced treadmill over 3-5 sessions (Figure 1A). After each habituation session, mice were returned to their home cage. Within one to two sessions, mice spontaneously locomoted in the forward direction interspersed with periods of rest or occasional sleep (Figure 1B). On the day of recording, mice were head-fixed. A 32- or 64-channel silicon probe/optrode was then advanced towards the STN, and electrophysiological recordings were made as mice rested and locomoted (Figure 1C). At the end of the recording session mice were perfuse-fixed. Recording sites were assessed histologically (Figure 1 D) and charted in 3-dimensions in the Allen Brain Atlas Common Coordinate Framework using NeuroInfo (MBF BioSciences).

**Figure 1.**
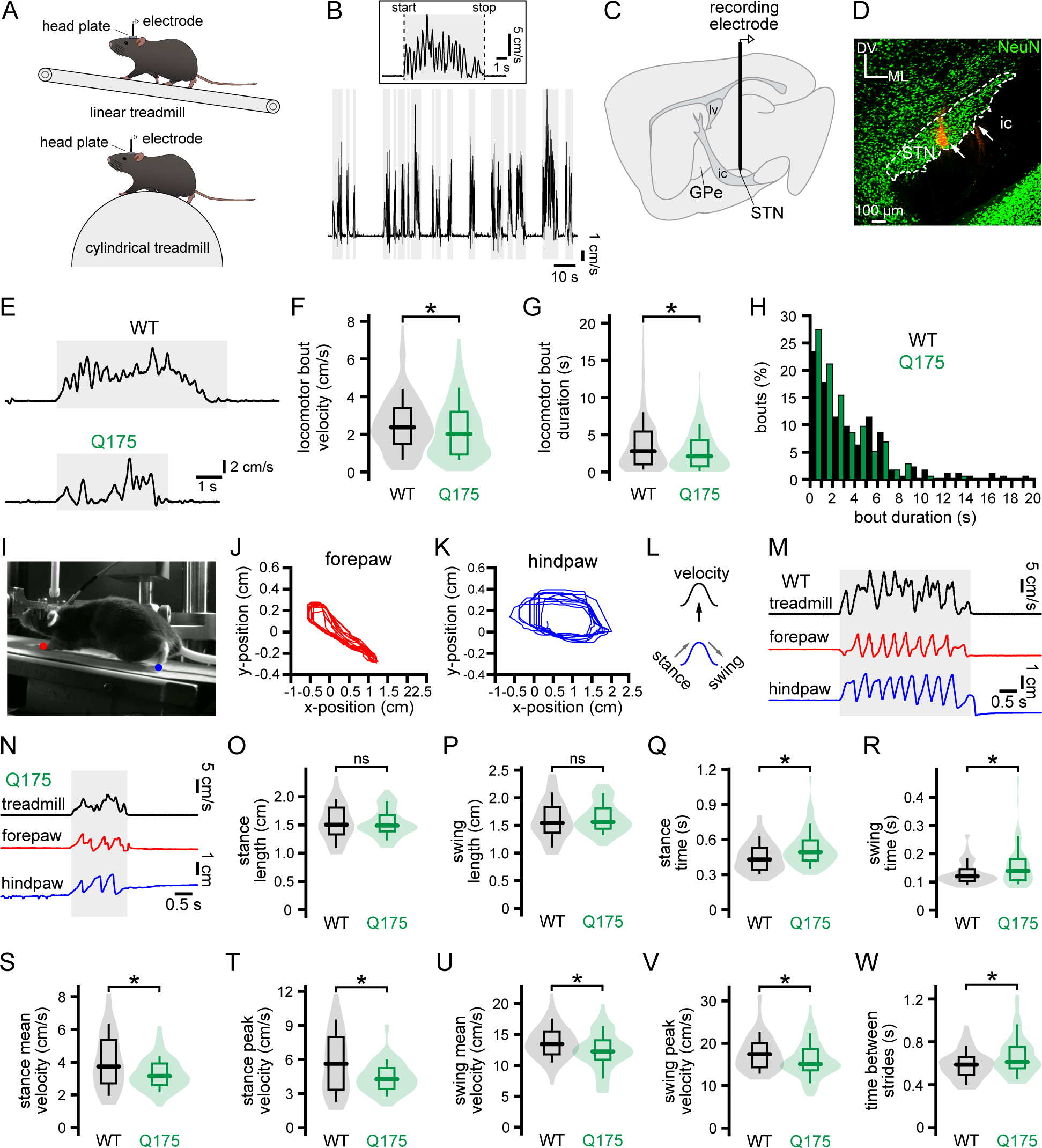
Self-initiated locomotion is dysregulated in Q175 HD mice. (A) STN encoding of self-initiated locomotion was assessed in head-fixed mice that were habituated to a self-paced linear or cylindrical treadmill. (B) A treadmill velocity encoder was used to detect periods of rest and bouts of self-initiated treadmill locomotion. (C) STN activity was recorded with a 32 or 64 channel silicon electrode/optrode. (D) The locations of STN recordings were assessed histologically using DiI red labeling (white arrows) and immunohistochemical detection of NeuN (green). The dorsoventral (DV) and mediolateral (ML) axes and internal capsule (ic) are denoted. In some cases, immunohistochemical detection of IBA1 was used to locate electrode tracks instead of DiI red (not illustrated). (E-H) Self-initiated treadmill locomotor bouts were shorter in duration and of lower velocity in Q175 HD mice versus WT mice (E, representative treadmill velocity encoder traces; F, G, H, population data). (I-N) DeepLabCut-based analysis of high frame rate digital video was used to track 2D contralateral paw movement in the x- and y-axes during self-initiated locomotion and rest (I, head-fixed mouse on a linear treadmill, with fore-(red) and hind-paw (blue) tracking denoted; J, K, x-y coordinates of the fore-(J, red) and hind-(K, blue) paws during locomotion; L-N, schematized (L) and example relationships of treadmill velocity and x-axis displacement of the contralateral paws during treadmill locomotion in WT (M) and Q175 (N) mice). (O-W) X-axis kinematics of the contralateral hindpaw in WT and Q175 mice during locomotion. The lengths of the stride and swing phases of locomotion were not significantly different in Q175 and WT mice (O, P). The durations (Q, R) and velocities (S-V) of both the stride and swing phases were longer and lower, respectively, in Q175 versus WT mice. During locomotion the time between strides was longer in Q175 mice (W). *, p < 0.05; ns, not significant.

Locomotor periods were defined as periods in which treadmill velocity ≥ 0.25 cm/s for ≥ 200 ms (Figure 1B). Pre- and post-locomotor periods were defined as the second preceding and following a locomotor bout, respectively. Self-initiated locomotion was of higher velocity in WT and Q175 female mice (Table S1) compared to their male counterparts. As a result, phenotypic comparisons were made using datasets that were matched for sex (4 males and 3 females per group). Although both WT and Q175 mice locomoted spontaneously in the head-fixed configuration, locomotor performance was subtly but consistently impaired in Q175 mice, analogous to deficits in freely moving mice^58–61^. Thus, locomotor bouts were of lower velocity and shorter duration (Figures 1E-H; Table S1). In some cases digital-movies (100 fps) of the mice contralateral to the recorded hemisphere were taken during the recording session and movements of the forepaw and hindpaw were tracked using DeepLabCut^63,64^ (Figures 1I-N). Because tracking was most accurate and consistent for the contralateral hindpaw, locomotion kinematics were assessed from this body part. Although, the contralateral hindpaw exhibited similar trajectories during locomotion in WT and Q175 mice (Figures 1O-P; Table S1), stance- and swing-associated movements were of longer duration (Figures 1Q, R; Table S1) and associated with lower mean and peak velocities (Figures 1S-V; Table S1) in Q175 mice. The interval between strides was also longer in Q175 mice (Figure 1W; Table S1).

To restrict our sample of STN activity to well-isolated single units, the following inclusion criteria were utilized: (1) PCA clusters were significantly different (p < 0.05); (2) J3-statistic ≥ 1; (3) Davies Bouldin test statistic ≤ 0.5. In addition, a threshold of < 0.5% of interspike intervals under 2 ms was required for classification as a putative single unit (% interspike interval within 2 ms; WT: 0.074, 0.0-0.2, n = 99; Q175: 0.0, 0.0-0.117, n = 103; values represent median and interquartile range). 30 second periods of immobility prior to pre-locomotor periods were used to measure STN activity at rest. At rest STN neurons in WT and Q175 mice discharged in a tonic but irregular firing pattern, as described previously^50^ (Figure 2A; Table S2). However, the overall frequency of STN activity was lower in Q175 mice (Figures 2A-C; Table S2), apparently due to a large increase in neurons discharging at frequencies below 5 Hz (Figure 2C; Table S2). The regularity of baseline activity, as assessed from the coefficient of variation (CV) of the interspike interval, was also significantly lower in Q175 than WT mice (Figures 2A and D-F; Table S2), including those STN neurons that fired below 5 Hz or at 5 Hz and above (Figure 2F; Table S2).

**Figure 2.**
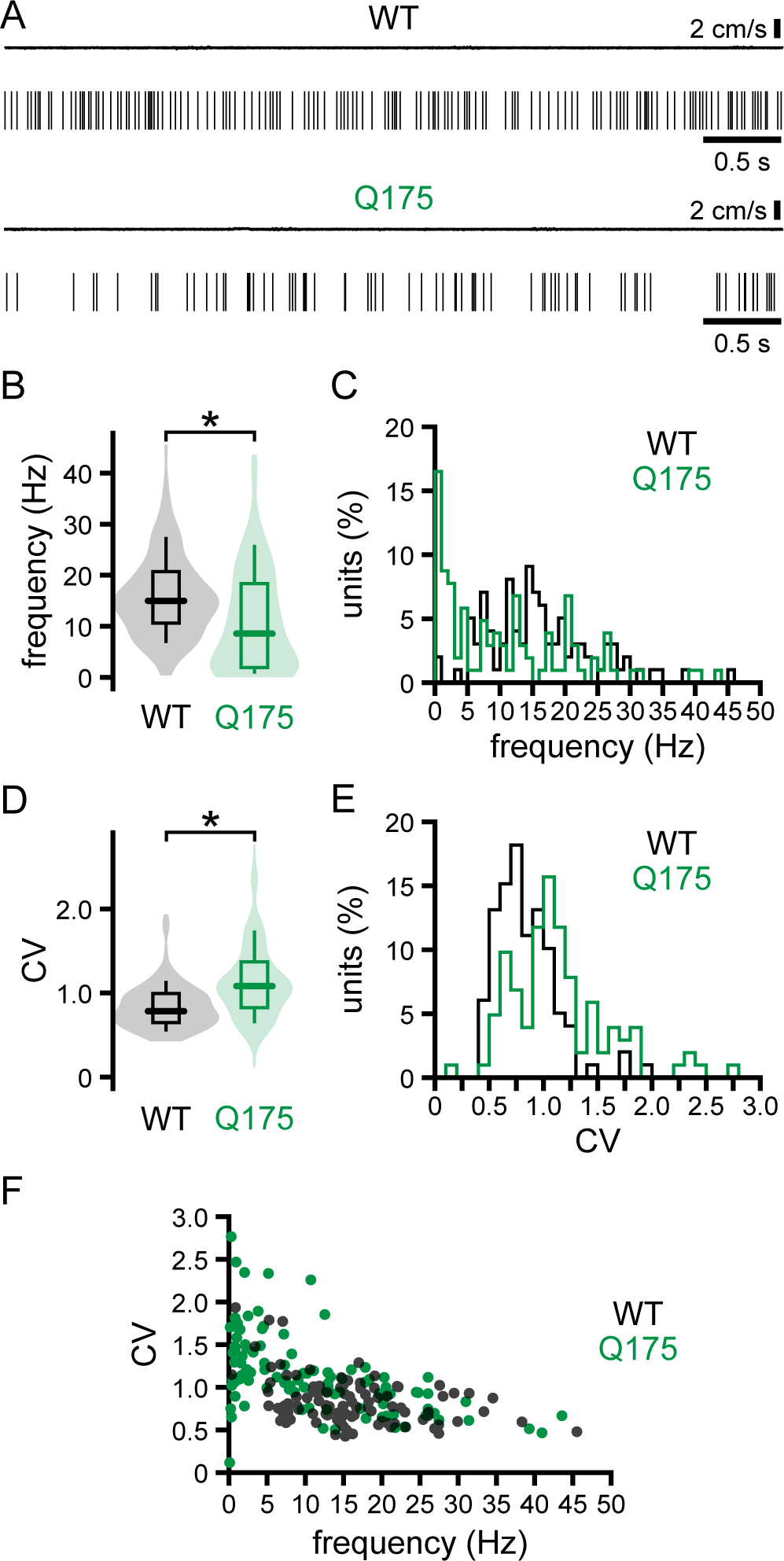
The frequency and precision of STN unit activity are lower in Q175 HD versus WT mice at rest. (A) Representative spike rasters of STN units and associated treadmill velocity encoder records in WT and Q175 mice at rest. (B-F, population data) The mean frequency and precision (coefficient of variation of the interspike interval, CV) of STN activity are lower in resting Q175 mice compared to WT. *, p < 0.05.

The majority of STN neurons in WT and Q175 mice consistently exhibited locomotion-related changes in firing that were positively correlated with treadmill velocity (WT: 64.9%, 61 of 94 neurons; Q175: 69.3%, 52 of 75 neurons; Figures 3A-N and S1A-B; Table S3). Units exhibiting these properties were defined as type 1 neurons. Overall, the firing frequency of type 1 units and their corresponding z-scores were greater in the pre-locomotion period versus rest, in the locomotion period versus the pre-locomotion and post-locomotion periods, and the post-locomotion period versus the subsequent rest period in both WT and Q175 mice (Figures 3F-G, M-N, and S1; Table S3). Spike frequency-treadmill velocity correlations ranged from weak to strong and although they generally peaked close to 0 ms, they were temporally broad in nature (Figures 3C, D, J, and K). Close inspection of instantaneous spike frequency versus treadmill velocity within a locomotor bout revealed that even for strongly correlated neurons, spike frequency varied widely for identical treadmill velocities and conversely could be similar for quite different treadmill velocities (Figures 3E and L). Together these data reveal that the majority of STN neurons exhibit locomotion-related increases in spiking activity that are positively correlated with treadmill velocity. However, the complexity of the instantaneous spike frequency-velocity relationships, and the modest and temporally broad nature of the spike frequency-velocity correlation argue that additional aspects of locomotion are encoded by type 1 cells.

**Figure 3.**
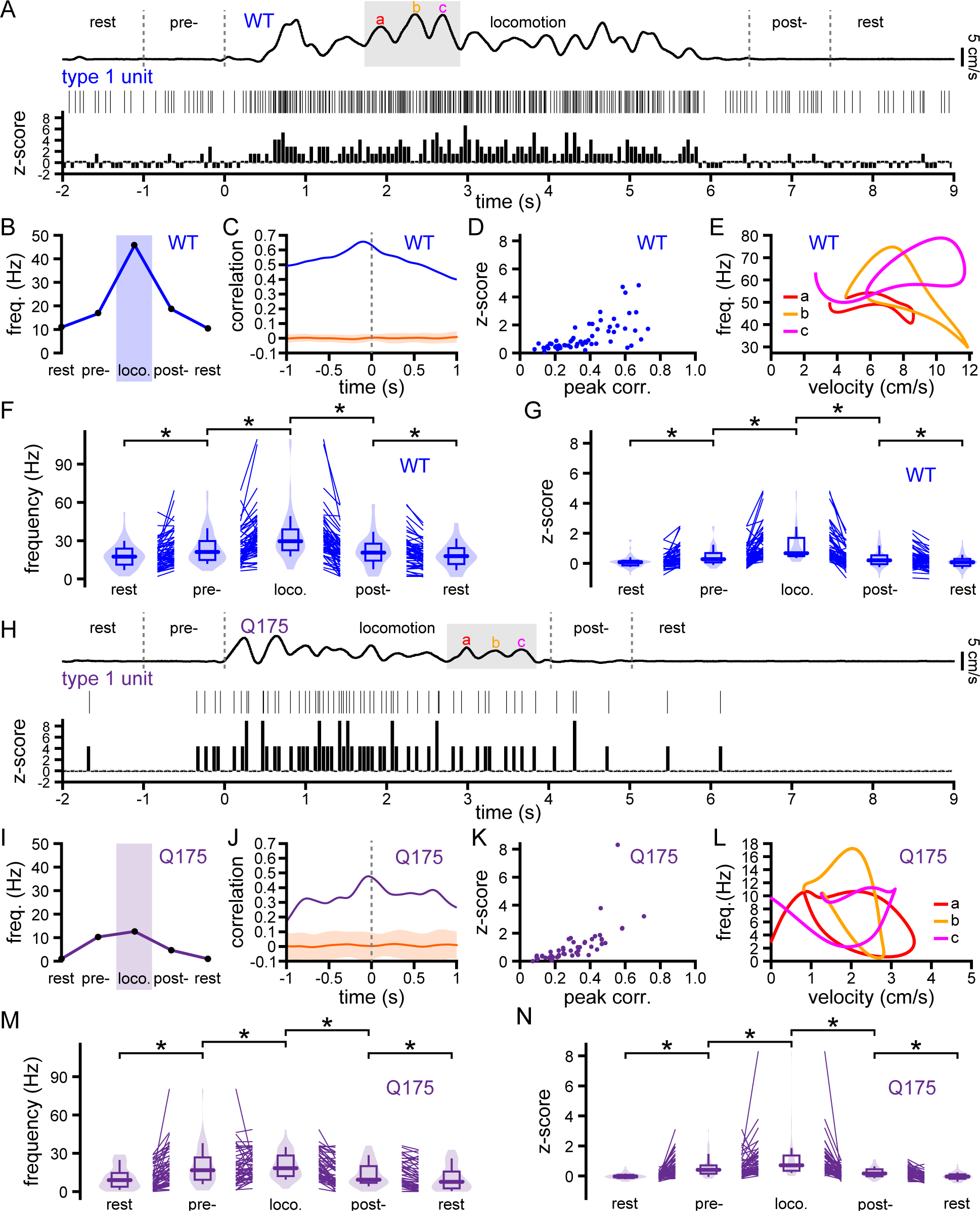
Type 1 STN neurons in WT and Q175 mice exhibit locomotion-associated increases in firing. (A) Locomotion-associated activity of a representative type 1 STN neuron in a WT mouse (upper trace, treadmill velocity; middle trace, spike raster; lower trace, z-score of spikes per 50 ms time bin relative to baseline spike counts). (B) Average firing frequency of the neuron in A during the initial rest, pre-locomotion, locomotion, post-locomotion, and subsequent rest periods. (C) Spike frequency-treadmill velocity correlation (blue) versus the correlation after shuffling (orange, mean +/- 2SD). (D) Peak correlation versus z-score for the sample population. (E) The spiking rate of the neuron in A varied inconsistently over several cycles of velocity change (a, b, c; color coded as for A). (F-G) Frequency (F) and z-score (G) population data for type 1 neurons in WT mice. (H) Locomotion-associated activity of a representative type 1 STN neuron in a Q175 mouse (upper trace, treadmill velocity; middle trace, spike raster; lower trace, z-score of spikes per 50 ms time bin relative to baseline spike counts). (I) Average firing frequency of the neuron in H during the initial rest, pre-locomotion, locomotion, post-locomotion, and subsequent rest periods. (J) Spike frequency-treadmill velocity correlation (purple) versus the correlation after shuffling (orange, mean +/- 2SD). (K) Peak correlation versus z-score for the sample population. (L) The spiking rate of the neuron in H varied inconsistently over several cycles of velocity change (a, b, c; color coded as for H). (M-N) Frequency (M) and z-score (N) population data for type 1 neurons in Q175 mice. *, p < 0.05.

Consistent with previous observations in anesthetized mice^50^, the mean frequencies of type 1 STN activity in the rest, pre-locomotor, locomotor, and post-locomotor periods were lower in Q175 mice (Figure 4A; Table S4). This was accompanied by a reduction in spike frequency-treadmill velocity correlation (WT: 0.3456, 0.2205-0.5145; Q175: 0.2837, 0.1779-0.4094; p = 0.0348; Table S4) in Q175 mice relative to WT. In contrast, the respective z-scores for each period were not significantly different in WT and Q175 mice (Figure 4B; Table S4). Together, these data argue that locomotion-related, synaptic patterning of type 1 neurons is similar in WT and Q175 mice but firing in response to synaptic input may be limited by the lower intrinsic excitability of STN neurons and hyperactivity of upstream prototypic GABAergic GPe neurons in Q175 mice, as described previously^50,52^. Although the frequency and CV of neuronal firing are often inversely related, spiking frequency and CV exhibited parallel locomotion related-changes in WT mice (Figure S1C). Thus, the synaptic mechanisms that underlie locomotion related increases in activity additionally confer irregularity in WT mice. This trend was less apparent in Q175 mice (Figure S1D). Overall, these data reveal that during rest and the pre-locomotor, locomotor, and post-locomotor periods, the frequency of type 1 unit activity is significantly lower in Q175 mice, although locomotion-related changes in activity are still present.

**Figure 4.**
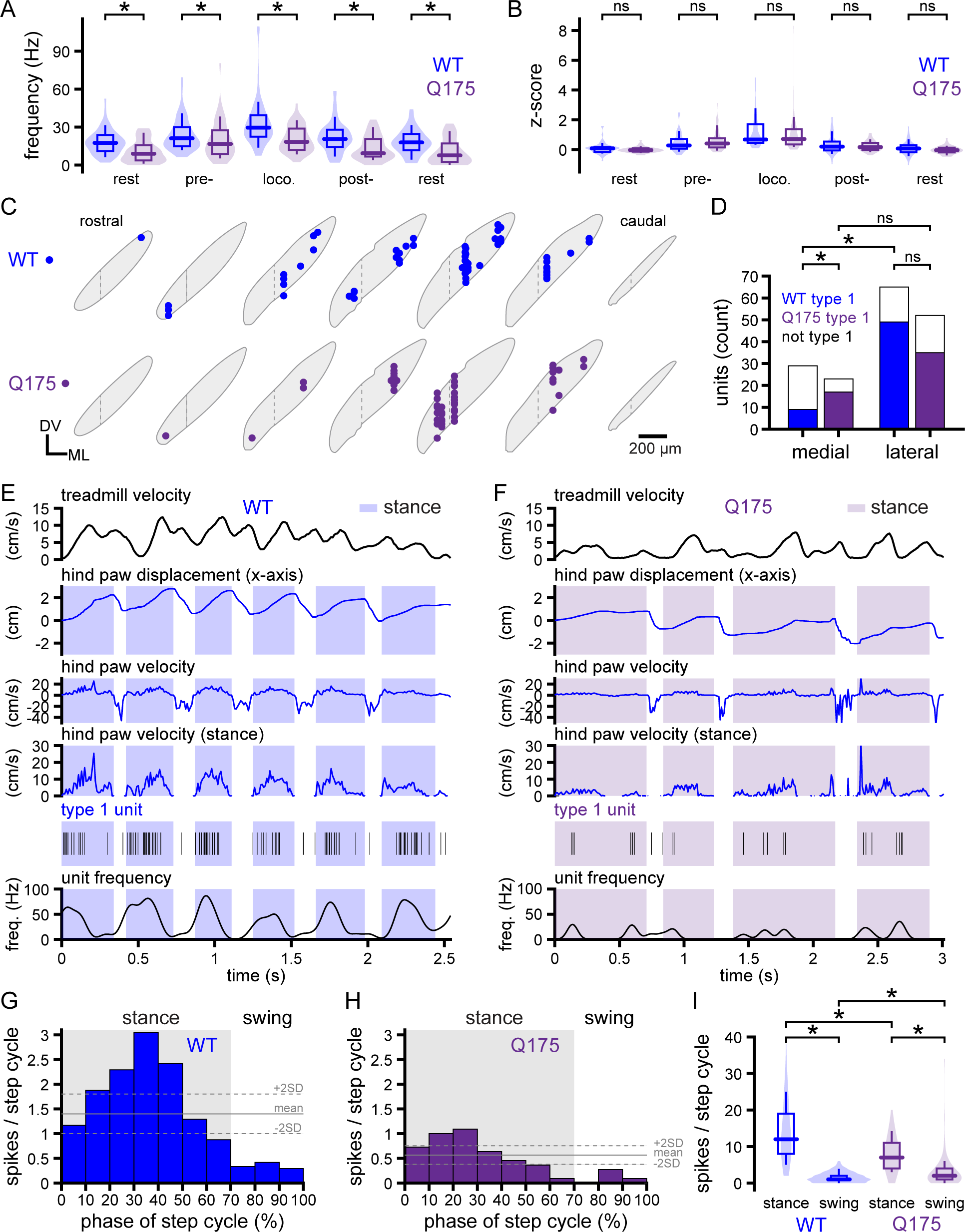
The frequencies of resting and locomotion-associated type 1 STN activity are reduced in Q175 mice. (A-B) Population data. The frequency (A) but not z-score (B) of locomotion-associated type 1 STN activity was reduced in Q175 mice. (C-D) Distribution of recorded type 1 STN neurons in WT and Q175 mice. In WT mice type 1 units were relatively abundant in the dorsolateral two-thirds of the STN compared to the medial third. In the medial third of the STN, type 1 units were more prevalent in Q175 than WT mice. The boundary between the medial third and lateral two-thirds of the STN is denoted by a dashed line. (E-I) A subset of type 1 STN neurons in WT and Q175 mice exhibited firing that was related to the phase of locomotor cycle (E-H, representative examples; I, population data). (I) The number of spikes during the stance phase of the locomotor cycle was significantly lower in Q175 mice. *, p < 0.05; ns, not significant.

Recordings in WT were slightly biased to the lateral two-thirds of the STN (74.1%) compared to the medial third (25.93%), in contrast to recordings in Q175 mice, which sampled these sectors as one would predict from the relative sizes of these domains, i.e. 66.7% of electrode tracks were in the lateral two-thirds of the STN and 33.3% were in the medial third (Figures S1E, F). Adjusting the numbers of type 1 units for sampling bias in this dimension did not alter the conclusion that the majority of STN units are type 1 in WT mice (type 1 = 61.7 %, 58 of 94 neurons). Furthermore, comparing the raw or adjusted numbers of type 1 units versus all other units revealed that type 1 units were concentrated in the lateral two-thirds of the STN in WT (WT raw: lateral STN type 1 = 54, not-type 1 = 17; medial STN type 1 = 7, not-type 1 = 16; p < 0.001, Fisher’s test; WT adjusted:lateral STN type 1 = 49, not-type 1 = 16; medial STN type 1 = 9, not-type 1 = 20; p < 0.001, Fisher’s test) but not Q175 mice (Q175 raw: lateral STN type 1 = 35, not-type 1 = 17; medial STN type 1 = 17, not-type 1 = 6; p > 0.05, Fisher’s test) (Figures 4C and D). Whether in Q175 mice this represents a loss of type 1 encoding in lateral STN neurons, or a developmental shift in the spatial distribution of type 1 neurons or their afferents is unclear.

The activity of a subset of type 1 units was further analyzed with respect to contralateral limb kinematics during locomotion. 50% and 35% of units in WT and Q175 mice, respectively, exhibited firing that was related to the phase of the locomotor cycle, as assessed from histograms of spikes versus phase of locomotion (WT: n = 6 of 12 neurons; Q175: n = 6 of 17 neurons; Figures 4E-I and S2A-B; Table S4). Activity was highest during the stance or propulsive phase of the contralateral hindpaw locomotor cycle (Figures 4E-I; Table S4). In neurons with phase-encoding, the total number of spikes per stance phase was lower in Q175 mice, reflecting the general hypoactivity of type 1 units in these mice (Figures 4E-I; Table S4). 50% and 65% of units in WT and Q175 mice, respectively, exhibited firing that was not related to the phase of the locomotor cycle (WT: n = 6 of 12 neurons; Q175: n = 11 of 17 neurons; Figures S2A-G; Table S4). In these cases, spike counts were similar during the stance/propulsive and swing phases of the contralateral hindpaw locomotor cycle, as assessed from histograms of spikes versus phase of locomotion(Figures S2A-G; Table S4). In neurons without phase-encoding, the number of spikes per swing phase was lower in Q175 mice (Figures S2C-G; Table S4).

In WT and Q175 mice, a minority of STN neurons (type 2) consistently exhibited firing rates that decreased during locomotion whether adjusted for sampling bias in WT or not (WT raw: 28%, n = 26 of 94; Q175: 16%,12 of 75; Figures 5, S3A and B; Table S5) and were negatively correlated with treadmill velocity (Figures 5C, D, J, and K; Table S5). In contrast to type 1 neurons, the frequencies of pre-locomotor activity were not significantly different from the preceding rest period (Figures 5F and M; Table S5). However, the frequency and associated z-scores, and precision of firing during locomotion were significantly lower than the pre-locomotor and post-locomotor periods (Figures 5F, G, M, N, and S3C, D). The firing rates and/or associated z-scores were also modestly but significantly elevated in the second following locomotion compared to the subsequent rest period (Figures 5F, G, M, and N) in WT and Q175 mice. If the locomotion-associated dips in type 2 activity are due to synaptic inhibition, the elevated firing in the post-locomotor period could reflect post-inhibitory rebound firing, an intrinsic membrane property of most STN neurons^65,66^. In stark contrast to type 1 units, the frequency and pattern of type 2 neuron activity at rest and during locomotion were not significantly different in WT and Q175 mice (Figures 6A, B, and S3C, D; Table S6). Type 2 units were relatively prevalent in the medial STN of WT compared to the lateral STN, but exhibited no spatial preference in Q175 mice, whether corrected for sampling bias or not (WT raw: lateral STN type 2 = 14, not-type 2 = 57; medial STN type 2 = 12, not-type 2 = 11; p = 0.030, Fisher’s test; WT adjusted: lateral STN type 2 = 13, not-type 2 = 52; medial STN type 2 = 15, not-type 2 = 14; p = 0.0156, Fisher’s test; Q175 raw: lateral STN type 2 = 9, not-type 1 = 43; medial STN type 2 = 3, not-type 2 = 20; p > 0.05, Fisher’s test) (Figures 6C and D). During locomotion the firing of type 2 neurons was not consistently related to the phase of the locomotor cycle and the numbers of spikes per stance and swing phase were similar in WT and Q175 mice (Figures 5E, L, and 6E-I); Tables S5 and S6). Together these data demonstrate that in contrast to type 1 neurons, the rate and pattern of type 2 neuron activity both at rest and during locomotion are similar in WT and Q175 mice. The firing of a small proportion of STN neurons was uncorrelated with locomotion (WT: 7%, n = 7 of 94 neurons; Q175: 15%, n = 11 of 75 neurons). The spatial distribution of these neurons did not exhibit a preference for the lateral or medial STN in WT or Q175 mice (Figure 6C).

**Figure 5.**
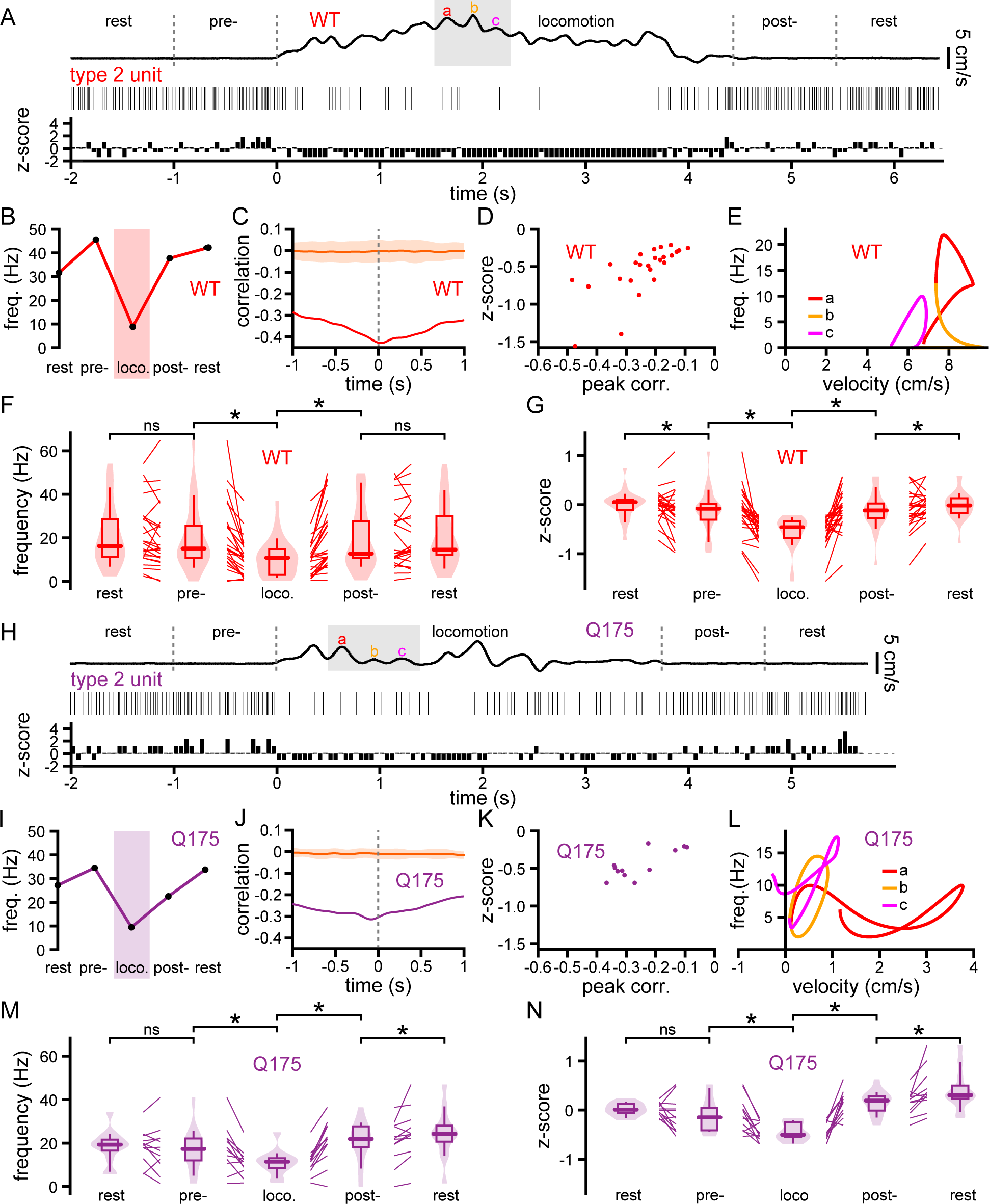
Type 2 STN neurons in WT and Q175 mice exhibit locomotion-associated decreases in firing. (A) Locomotion-associated activity of a representative type 2 STN neuron in a WT mouse (upper trace, treadmill velocity; middle trace, spike raster; lower trace, z-score of spikes per 50 ms time bin relative to baseline spike counts). (B) Average firing frequency of the neuron in A during the initial rest, pre-locomotion, locomotion, post-locomotion, and subsequent rest periods. (C) Spike frequency-treadmill velocity correlation (red) versus the correlation after shuffling (orange, mean +/- 2SD). (D) Peak correlation versus z-score for the sample population. (E) The spiking rate of the neuron in A varied inconsistently over several cycles of velocity change (a, b, c; color coded as for A). (F-G) Frequency (F) and z-score (G) population data for type 2 neurons in WT mice. (H) Locomotion-associated activity of a representative type 2 STN neuron in a Q175 mouse (upper trace, treadmill velocity; middle trace, spike raster; lower trace, z-score of spikes per 50 ms time bin relative to baseline spike counts). (I) Average firing frequency of the neuron in H during the initial rest, pre-locomotion, locomotion, post-locomotion, and subsequent rest periods. (J) Spike frequency-treadmill velocity correlation (purple) versus the correlation after shuffling (orange, mean +/- 2SD). (K) Peak correlation versus z-score for the sample population. (L) The spiking rate of the neuron in H varied inconsistently over several cycles of velocity change (a, b, c in H are color coded). (M-N) Frequency (M) and z-score (N) population data for type 2 neurons in Q175 mice. *, p < 0.05; ns, not significant.

**Figure 6.**
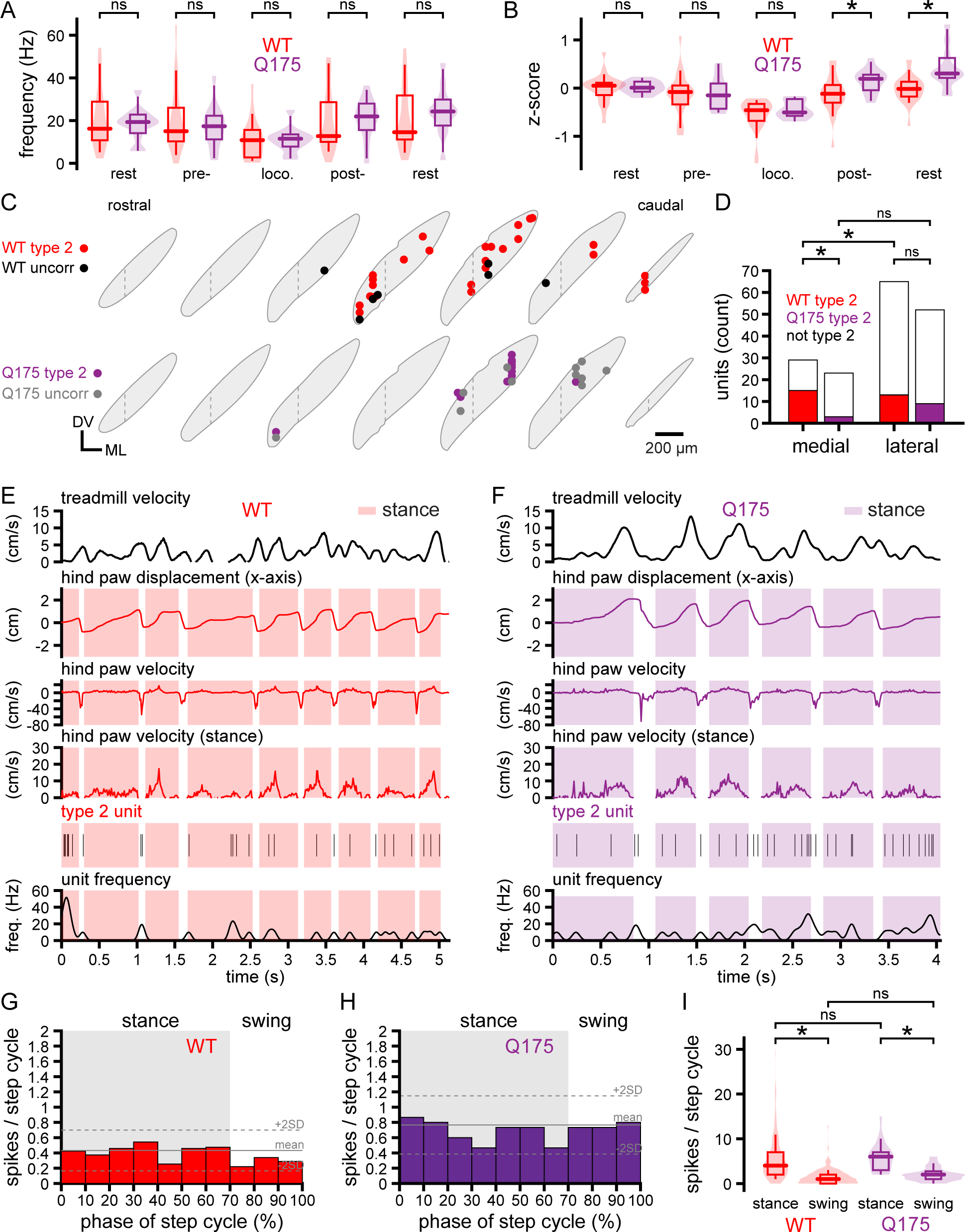
The frequencies and patterns of resting and locomotion-associated type 2 STN neuron activity are similar in WT and Q175 HD mice. (A-B) Population data. The frequencies (A) and z-scores (B) of resting and locomotion-associated type 2 STN activity were similar in WT and Q175 mice. (C-D) Distribution of recorded type 2 STN neurons in WT and Q175 mice. The boundary between the medial third and lateral two-thirds of the STN is denoted by a dashed line. Type 2 units were located throughout the dorsolateral extent of the caudal half of the STN in WT and Q175 mice. Type 2 units were 1) more abundant in the medial than the lateral STN of WT mice 2) more abundant in the medial STN of WT mice than the medial STN of Q175 mice. The distribution of units whose activity was uncorrelated with locomotion are also Illustrated (C). (E-I) Type 2 STN neurons in WT and Q175 mice exhibited firing that was unrelated to the phase of the locomotor cycle (E-H, representative examples; I, population data). (I) The number of spikes per locomotor cycle was similar in WT and Q175 mice. *, p < 0.05; ns, not significant.

Several studies have revealed that the molecular properties of STN neurons are heterogeneous, e.g., subtypes of STN neurons differentially express the calcium binding protein parvalbumin (PV) or the type 2 vesicular glutamate transporter (vGluT2) and exhibit distinct membrane and synaptic properties^44–46^. To determine whether molecularly defined STN subtypes are correlated with STN subtypes defined by their locomotion-encoding properties, STN subtypes were optogenetically tagged. The encoding properties and non-specific effects of light delivery were first established in control WT mice in which eGFP was virally expressed in STN neurons through injection of AAVs carrying a synapsin promoter-dependent construct (Figures S4A, B, F, G, H, K, and S5A, B; Table S7). In the absence of opsin expression, 633 nm light delivery had no consistent effect on the activity of 11 units that were recorded (Figures S4F, G, and S5A; Table S7). Of these 11 units, 8 exhibited type 1 activity, 2 exhibited type 2 activity, and 1 was uncorrelated with locomotion (Figures S4H, K, and S5B; Table S7), similar to the proportions of encoding subtypes described above.

To determine whether PV expression is correlated with the locomotion-encoding subtypes described above, eNpHR3.0-eYFP or ChR2(H134R)-eYFP was expressed in PV+ STN neurons through injection of AAV vectors carrying Cre-dependent constructs into the STN of PV-Cre mice (Figures S4A, C, D, F, G, I, L and S5C, D, E, F; Table S7). Silicon optrodes were then used to optotag/identify and record the activity of STN PV+ neurons at rest and during locomotion (Figures S4A, C, D, F, G, I, L and S5C, D, E, F; Table S7). Using this approach 11 neurons were optotagged. Of these 9 exhibited type 1 neuronal activity and 2 exhibited type 2 neuronal activity (Figures S4I, L and S5F; Table S7). A similar approach was used to determine whether the subset of STN neurons that express vGluT2 exhibit distinct encoding properties^46^. In this case the inhibitory opsin eNpHR3.0-eYFP was virally expressed in a Cre-dependent manner in vGluT2-Cre mice (Figures S4A, E, F, G, J, M and S5G, H; Table S7). Using this approach, 11 neurons were optotagged. Of these 7 exhibited type 1 neuronal activity and 3 exhibited type 2 neuronal activity and 1 exhibited activity that was uncorrelated with locomotion (Figures S4J, M and S5H; Table S7). Given that the proportions and distributions of optotagged STN neurons exhibiting type 1 or type 2 neuronal activity were similar to neurons expressing eGFP (Figures S5I and J; Table S7) or non-expressing neurons described above, we conclude that neither PV or vGluT2 expression reliably distinguishes between STN neurons with type 1 or 2 locomotor encoding properties. The baseline and locomotion-associated activities and locomotion-associated z-scores of identified PV+ and vGluT2+ STN neurons were also not significantly different from each other or eGFP-expressing STN neurons (Figures S5K, L, and M; Table S7).

To determine the functional role of locomotion-associated STN activity, the inhibitory opsin eNpHR3.0-eYFP was virally expressed in vGluT2-Cre mice through subthalamic injection of AAVs carrying a Cre-dependent construct, as described above (Figures S4A and E), or in WT mice through subthalamic injection of AAVs carrying a CaMKII-dependent expression construct (Figures S6A-D). To control for the effects of viral expression and light delivery, eGFP alone was expressed in the STN in a subset of WT mice, as described above (Figure S4A, B). 32-64 channel silicon optrodes were then used to record and unilaterally inhibit STN activity for 5-seconds in mice at rest or during a self-initiated locomotor bout (Figures 7 and S6; Table S8). Data from WT or vGluT2-Cre mice were pooled because activation of eNpHR3.0-eYFP inhibited STN activity similarly (Figure S6E; Table S8). Application of 633 nm light in eGFP-expressing WT mice had no effect on STN unit activity, as described above (Figures S4F, G; Tables S7 and S8). Consistent with the majority of neurons in the lateral two-thirds of the STN being type 1 (Tables S4 and S6), optogenetically inhibited neurons in this sector were predominantly type 1 encoding (type 1, 90.9%; type 2 9.1%; uncorrelated, 0%; Figure S6F; Table S8). In contrast, in the medial third of the STN where encoding subtypes are more evenly represented (Figure S6F; Tables S4 and S6), optogenetically inhibited STN neurons were a mixture of subtypes (type 1, 27.3%; type 2, 45.4%; uncorrelated, 27.3%; Table S8). Optogenetic inhibition of the lateral two-thirds or medial third of the STN or distinct encoding subtypes suppressed unit activity to a similar degree (Figures 7A, B; Table S8).

**Figure 7.**
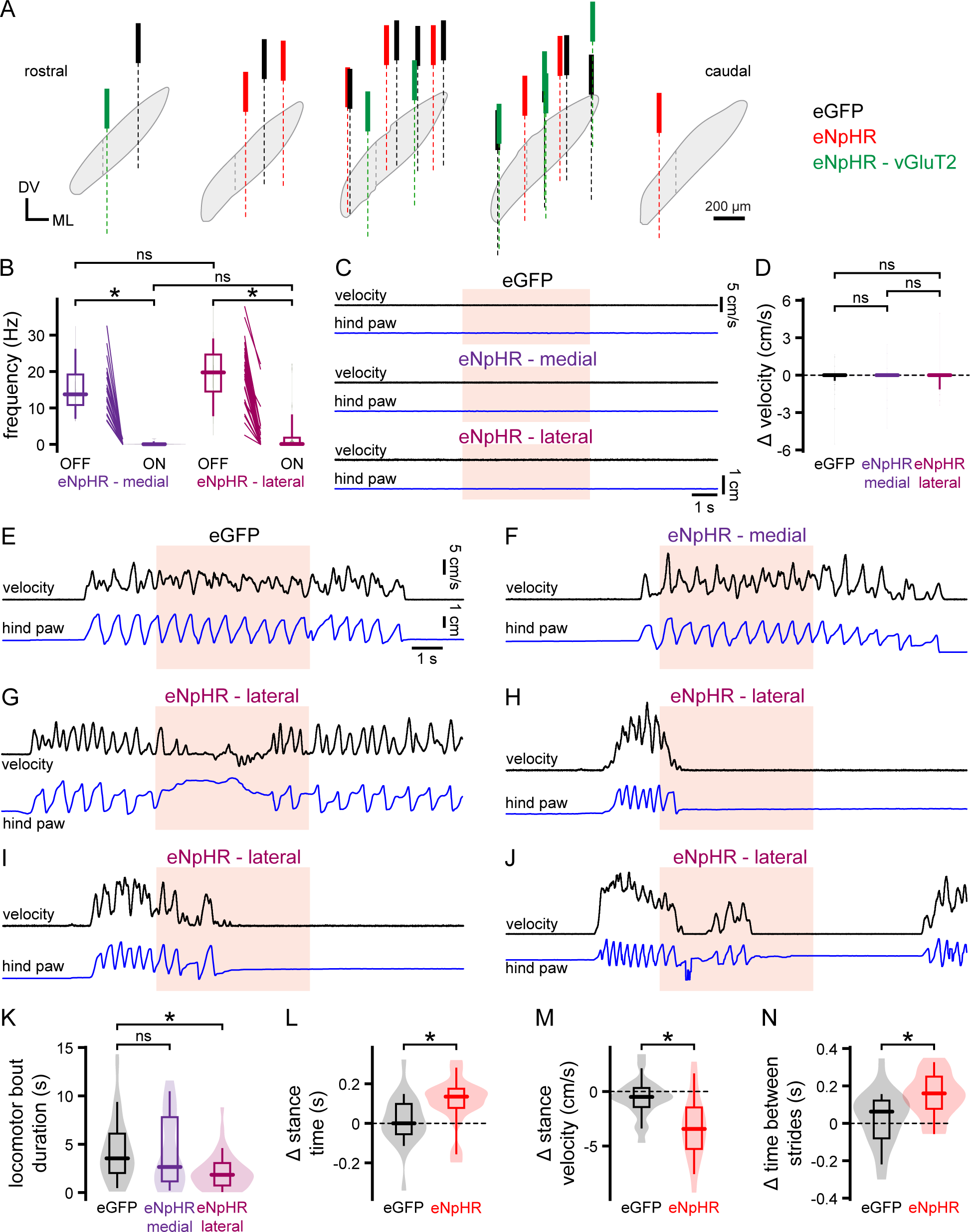
Optogenetic inhibition of movement-related STN activity dysregulates locomotion. (A) Positions of the fiberoptics used to deliver 633 nm light to the STN in control eGFP-expressing WT mice (black), NpHR3.0-eYFP-expressing WT mice (red), and NpHR3.0-eYFP-expressing vGluT2-Cre mice (green). The boundary between the medial third and lateral two-thirds of the STN is denoted by a dashed line. (B) Impact of stimulating eNpHR3.0-eYFP in the medial third or lateral two-thirds of the STN on neuronal activity. (C-D) Effects of 633 nm STN light delivery on eGFP- or NpHR3.0-eYFP-expressing resting mice on treadmill velocity and hindpaw kinematics (C, representative examples; upper trace, treadmill velocity; lower trace, hindpaw x-axis displacement; D, population data). (E-J) Representative examples of the effects of 633 nm light delivery on eGFP- or NpHR3.0-eYFP-expressing mice during locomotion (upper traces, treadmill velocity; lower traces, hindpaw x-axis displacement). Relative to the effects of light delivery in eGFP-expressing control mice (E) optogenetic inhibition of the medial STN had no effect on locomotion (F). In contrast, optogenetic inhibition of the lateral STN dysregulated, slowed, and more rapidly terminated locomotion compared to locomotor bouts in control mice (G-J). (K-L) Population data. Compared to the effects of 633 nm light in eGFP-expressing control mice, optogenetic inhibition of the dorsolateral STN reduced the duration of self-initiated locomotor bouts (K; optogenetic inhibition of the medial STN had no significant effect on bout duration relative to control). Based on contralateral hindpaw kinematics, optogenetic inhibition of the lateral STN also increased the duration (L), reduced the velocity (M) of the stance phase, and increased the time between strides (N). *, p < 0.05; ns, not significant.

Optogenetic inhibition of the lateral or medial STN or equivalent light delivery in eGFP-expressing control mice had no effect at rest arguing that reduction of type 2 STN activity is not sufficient for locomotion initiation, (Figures 7C and D; Table S8). Optogenetic inhibition of the medial third of the STN also had no significant effect on coincident locomotion relative to the effects of identical light delivery in eGFP-expressing control mice (Figures 7E, F; Table S8). However, optogenetic inhibition of the lateral two-thirds of the STN interrupted coincident locomotion compared to the effects of similarly directed, timed, and intensities of light in eGFP-expressing mice (Figures 7E-N; Table S8). Optogenetic truncation of locomotion was associated with either abrupt time-locked cessation of locomotion, progressive slowing of locomotion before bout termination, and in some cases dysregulated limb movement, including reverse locomotion (Figures 7G-K; Table S8).

Given the small size of the STN and the potential for light to spread in the mediolateral axis, the regional effects of optogenetic inhibition were further analyzed. Optogenetic inhibition with fiberoptics placed over the lateral or central third of the STN truncated locomotor bouts to a similar degree versus optogenetic inhibition with fiberoptics over the medial third of the STN or light delivery alone (Figure S6G, Table S8). Together these data confirm that only optogenetic inhibition of the lateral two-thirds of the STN disrupts locomotion. Optogenetic inhibition similarly truncated locomotion in WT and vGluT2-Cre mice (Figure S6H; Table 8) and did not have cumulative effects on locomotion (Figure S6I; Table S8) because locomotor bouts that coincided with optogenetic inhibition in the first 50% or subsequent 50% of trials were of similar duration. Optogenetic inhibition of the lateral two-thirds of the STN also significantly disrupted contralateral hindpaw kinematics (Figures 7L-N; Table S8). Thus, the duration of the stance phase of the locomotor cycle (Figure 7L; Table S8) increased, the velocity of the stance phase decreased (Figure 7M; Table S8), and the interval between strides increased (Figure 7N; Table S8). Together, these data argue that 1) locomotion-locked type 1 unit activity in the lateral STN contributes to optimal locomotor performance 2) optogenetic inhibition of the lateral STN induces locomotor effects analogous to the deficits seen in Q175 mice, in which the activity of type 1 STN neurons both at rest and during locomotion are reduced relative to WT.

## Discussion

Similar to studies of arm movement-related subthalamic activity in primates^8,10,13,24,26^, in mice the majority of STN neurons (type 1) exhibited locomotion-dependent increases in activity. Although type 1 units (by definition) exhibited significant spike rate/treadmill velocity correlations, associated Pearson’s coefficients ranged from weak (0.1) to strong (0.7), presumably because the relationships between spike rate and treadmill velocity *within* a stride cycle or locomotor bout were inconsistent with units exhibiting very different spike rates for identical velocities. Furthermore, although spike rate/velocity correlations peak close to 0 ms, velocity encoding was temporally imprecise because correlation strength decayed modestly over a second. Together, these data argue that type 1 STN neurons encode additional aspects of volitional movement. Consistent with this possibility 1) 50% of type 1 neurons exhibited activity that was highest during the propulsive phase of the contralateral locomotor cycle 2) arm movement-related changes in STN discharge exhibit direction selectivity and/or correlate with movement amplitude/velocity in monkeys^14,24,26,27^ 3) changes in STN beta band activity are correlated with the onset, execution, vigor, phase, and termination of bipedal locomotion in human PD patients^28,30^. Finally, type I neurons may also encode movements that are less directly linked to locomotion such as axial postural adjustments, and/or micromovements unrelated to locomotion itself^67^.

Type 1 units typically exhibited increases in activity several hundred milliseconds prior to the onset of locomotion, consistent with several studies that detected elevations in STN activity prior to arm movement and associated changes in EMG^14,26^. These data suggest that type 1 activity is driven at least in part by motor command signals emanating from motor cortical regions. Following the end of each locomotor bout, type 1 activity decreased in a time-locked fashion. However, activity remained modestly elevated during the second following locomotion. This persistent elevation of firing may reflect ongoing activity in upstream nuclei related to post-locomotion postural adjustments and/or the slow decay of metabotropic receptor-mediated afferent synaptic transmission.

In concordance with previous studies and the distribution of cortical inputs, type 1 units were concentrated in the lateral two-thirds of the STN in WT mice. The circuits driving locomotion-related type 1 activity are potentially complex because the STN receives inputs from motor command structures such as the primary motor cortex and mesencephalic locomotor region and sensory/proprioceptive information via the cortex, parafascicular thalamic nucleus, and brainstem^30,68–71^. Indeed, neurons in the lateral STN respond to both motor cortical stimulation with short latency excitation, consistent with monosynaptic driving^13,23,38^, and to passive limb movement with increased activity, consistent with proprioceptive encoding^8,24,26,30^. It is also likely that indirect pathway-mediated striatopallidal inhibition of prototypic external globus pallidus (GPe) neurons leads to disinhibition of STN type 1 neurons during locomotion^23,38,72,73^. Indeed, neurons in the lateral STN also respond to motor cortical stimulation with a second longer latency and duration excitation due to indirect pathway-mediated disinhibition^23,38,39^. A recent study suggests that parafascicular thalamic inputs to the STN can also potently drive locomotion under some conditions^71^.

Approximately 28% of STN neurons consistently exhibited locomotion-locked reductions in activity across locomotor bouts in WT mice. In contrast to type 1 units, changes in type 2 activity were restricted to the locomotor period only. Type 2 units exhibited negative weak to strong spike rate/treadmill velocity correlations that were temporally broad. As for type 1 neurons, the relationships between spike rate and treadmill velocity within a locomotor bout were inconsistent. However, the locomotor cycle was not well represented by type 2 activity, presumably due to its sparsity. Together these data argue that reductions/cessations in type 2 activity encode the period rather than the parameters of locomotion. Type 2 units were evenly distributed across the mediolateral axis of the STN and were often recorded concurrently with type 1 units. STN units with movement-related reductions in firing have also been reported for upper limb movement in primates and were similarly rare^13,24,26^. One possibility is that type 2 neurons encode the movement of body parts unrelated to locomotion and their activity is suppressed by a combination of low motor command/proprioceptive drive and/or increased inhibition from disinhibited prototypic GPe neurons that are not targets of striatopallidal inhibition^72,74,75^. Another possibility is that some of these neurons, particularly in the medial third of the nucleus, signal the stopping or switching of actions and are suppressed by low drive from stop/switch-encoding cortical areas (e.g., dorsomedial prefrontal cortex) and/or increased inhibition from disinhibited prototypic GPe neurons that are not subject to striatopallidal inhibition^9,36^. The relatively large proportion of STN neurons exhibiting locomotion-related activity in WT mice compared to contralateral arm-related activity in primates^8,10,24,26,34^ may reflect the involvement of all four limbs in quadrupedal locomotion and a lower degree of somatotopic specificity in rodents^23,31,35^. Less than 10% of STN units were uncorrelated with locomotion. Given their lack of engagement in movement-encoding, these neurons may exclusively convey stop or switch signals from cortical areas such as the dorsomedial prefrontal cortex^8,10,12,23,38,39^.

Multiple studies have demonstrated that STN neurons are heterogeneous in their molecular properties^44–46^. In many brain nuclei, this molecular diversity delineates cell classes that are embedded in distinct circuits that subserve specific functions^76^. To address this possibility, we optoagged PV- or vGluT2-expressing neurons and measured their activity at rest and during locomotion. Although these neuron classes exhibit distinct membrane and synaptic properties, and spatial distributions, and subserve distinct functions in some contexts^46^, PV- and vGluT2-expressing neurons were comprised of both type 1 and 2 STN neurons. Thus, PV and vGluT2 expression in the STN are unreliable markers of movement encoding and presumably confer molecular specializations related to other aspects of circuit function e.g., motor learning^46^.

Q175 and other HD mouse models exhibit progressive movement and gait deficits that mimic those seen in human patients, e.g., reductions in locomotion velocity and distance, stance and swing velocity, and stride coordination, and increases in inter-stride interval^59–61^. Consistent with these deficits, head-fixed Q175 mice exhibited relatively short durations and low velocities of locomotion, low velocity paw kinematics, and increased time between strides. These deficits were specifically associated with abnormal hypoactivity of type 1 STN neurons both at rest and during locomotion. Previous studies utilizing *ex vivo* brain slices from Q175 mice or anesthetized Q175 mice revealed that a subset of STN neurons exhibit hypoactivity relative to WT mice^52,77^. This hypoactivity was caused by 1) NMDA receptor-dependent mitochondrial oxidant stress and increased activation of K_ATP_ channels by reactive oxygen species (ROS)^52^ 2) hyperactivity of upstream GABAergic GPe neurons^77^. Given that locomotion-associated increases in STN activity could arise from increased glutamatergic drive from motor command and proprioceptive structures and disinhibitory signaling from the GPe, the hypoactivity of type 1 neurons in Q175 mice may reflect increased NMDA receptor and ROS-dependent activation of K_ATP_ channels *and* pathological hyperactivity of upstream GPe neurons. Consistent with these mechanisms, 1) mitochondrial oxidant stress, ROS generation, and K_ATP_ channel activation are elevated in *ex vivo* brain slices from Q175 mice^52^ 2) although optogenetic inhibition of prototypic GPe neurons disinhibited STN activity in both anesthetized WT and Q175 mice, STN activity remained lower in Q175 mice^77^.

To determine the role of the STN in regulating locomotor activity, we optogenetically inhibited the STN of WT mice for 5 seconds at rest or during locomotion. No effect of optogenetic inhibition was observed in resting mice, arguing that STN hypoactivity generally and cessation of type 2 neuron activity specifically are not sufficient to initiate locomotion. At first sight, this observation appears contrary to the effects of STN lesions or prolonged pharmacological/optogenetic/chemogenetic inhibition of the STN^5,13,15–18^. However, the dysregulated movement that accompanies STN lesions or prolonged subthalamic inhibition may require volitional motor commands for its expression. If correct, the short duration of the optogenetic inhibition utilized here may have sufficiently reduced the probability of coincident action initiation/execution that the behavioral impact of STN inhibition was not revealed. Consistent with this hypothesis, brief optogenetic inhibition of the STN rapidly dysregulated self-initiated locomotor bouts that were already in the process of execution. Importantly this effect was restricted to the inhibition of the lateral STN where type 1 units are concentrated. Analogous dysregulation of a highly trained action sequence or spontaneous behavior accompanies bidirectional optogenetic manipulation of upstream components of the indirect pathway/network^78^ and is in line with reports that concomitant, coordinated direct *and* indirect pathway activity is necessary for optimal volitional movement^79–81^. Thus, rather than simply stopping movement^82^, indirect pathway nuclei, including the STN, play key roles in movement execution.

Precisely how type 1 STN activity optimizes self-initiated locomotion remains to be tested. One possibility is that widespread inhibition of movement-generation/regulation circuits by the indirect and hyperdirect pathways that flow through the STN enhances the functional impact of selective direct pathway-mediated disinhibition of locomotion-generating circuitry^4,13,83^. That said, the richness of locomotor cycle encoding by 50% of type 1 neurons argues that the indirect and hyperdirect pathways may also play a more dynamic and active role in kinematic regulation than previously supposed^84^ e.g., by terminating direct pathway-mediated inhibition of basal ganglia output neurons^13^. Another possibility is that type 1 STN activity promotes locomotion through excitation of dorsal striatum-projecting substantia nigra (SN) dopamine neurons which in turn mediate D1 dopamine receptor-dependent enhancement of direct pathway striatal projection neuron excitability and synaptic integration^21,85–87^, but see^88–90^. However, optogenetic inhibition of SN dopamine neurons does not dysregulate motor sequences already in the process of execution^89^, arguing that the rapid impact of STN inhibition on locomotion is likely unrelated to interruption of the STN’s excitation of dopamine neurons.

How do the findings described here inform our understanding and treatment of movement disorders? Together with other studies, our data support the conclusion that type 1 STN activity helps to optimize gait and that dysregulation of this activity in HD or PD contribute to gait deficits in these movement disorders. In HD, the cell-autonomous effects of mutant huntingtin directly compromise the encoding and survival of a subset of STN neurons^52,62,77^, arguing that treatments that suppress mutant huntingtin expression^91^ should be targeted widely and not focused solely on the striatum or cortex for full circuit/functional rescue. In PD, incomplete restoration of subthalamic locomotor encoding may contribute to the mixed effects of repetitive high-frequency DBS of the STN on gait deficits^92^. Indeed, coordinated reset stimulation^93^ or locomotor cycle-dependent stimulation^28^ that restore or impose more natural patterns of STN activity, respectively, may have the potential to rescue gait more consistently than traditional STN DBS.

## Limitations of the study

The major limitation of this work is the absence of genetic tools that could be used to distinguish the functional roles of type 1 and 2 STN neurons. To circumvent this, optogenetic inhibition was directed to sectors of the STN where type 1 units were the major cell type (lateral STN) or type 2 neurons comprised the majority (medial STN). Although optogenetic inhibition of type 2 neurons was partly occluded by locomotion-associated suppression, optogenetic inhibition suppressed type 2 activity completely. Thus, optogenetic inhibition of type 2 activity in the lateral STN may have contributed to locomotor dysregulation. Similarly, optogenetic suppression of type 1 activity in WT mice was more severe than the hypoactivity of these neurons in Q175 mice. Another limitation of our work is that the circuit elements responsible for the generation of movement-related STN activity remain to be determined.

## Acknowledgements

This work was funded by grants from CHDI Foundation (A-5071), Aligning Science Across Parkinson’s (ASAP-020600), and NIH-NINDS (R01 NS041280 and R01 NS121174). The authors thank Sasha Ulrich, Danielle Rae Schowalter, and Marisha Alicea for maintenance of mouse colonies and Drs. Vahri Beaumont, Roger Cachope, and Ignacio Munoz-Sanjuan for helpful discussions and advice throughout the execution of this study.

## Author contributions

Conceptualization: JWC, MDB; Methodology: JWC, JCM, JFA, MDB; Validation: JWC, JCM, JFA, MDB; Formal Analysis: JWC, JCM, JFA, MDB; Investigation: JWC, DW, SK; Data Curation: JWC, MDB; Writing – Original Draft, JWC, MDB; Writing – Editing, JWC, JCM, JFA, DW, SK, MDB; Visualization, JWC, JCM, JFA, MDB; Supervision, JWC, MDB; Project Administration, JWC, MDB; Funding Acquisition, MDB.

## STAR METHODS

## KEY RESOURCES TABLE

| REAGENT or RESOURCE | SOURCE | IDENTIFIER |
| --- | --- | --- |
| **Antibodies** |  |  |
| Anti-Iba1, Rabbit antibody | FUJIFILM Wako Pure Chemical Corporation | RRID:AB_839504 |
| Anti-NeuN, Mouse antibody | Abcam | RRID:AB_10711040 |
| Anti-Parvalbumin, Guinea pig antibody | Synaptic Systems | RRID:AB_2156476 |
| Alexa Fluor 405, Donkey anti-Mouse IgG (H+L) | Thermo Fisher Scientific | RRID:AB_2884884 |
| Alexa Fluor 594-AffiniPure Donkey Anti-Guinea Pig IgG (H+L) | Jackson ImmunoResearch Laboratories, Inc. | RRID:AB_2340474 |
| Alexa Fluor 488-AffiniPure Donkey Anti-Mouse IgG (H+L) | Jackson ImmunoResearch Laboratories, Inc. | RRID:AB_2340846 |
| Alexa Fluor 594-AffiniPure Donkey Anti-Mouse IgG (H+L) | Jackson ImmunoResearch Laboratories, Inc. | RRID:AB_2340854 |
| Alexa Fluor 647-AffiniPure Donkey Anti-Mouse IgG (H+L) | Jackson ImmunoResearch Laboratories, Inc. | RRID:AB_2340863 |
| Alexa Fluor 488-AffiniPure Donkey Anti-Rabbit IgG (H+L) | Jackson ImmunoResearch Laboratories, Inc. | RRID:AB_2313584 |
| Alexa Fluor 594 AffiniPure Donkey Anti-Rabbit IgG (H+L) | Jackson ImmunoResearch Laboratories, Inc. | RRID:AB_2340621 |
| Alexa Fluor 647-AffiniPure Donkey Anti-Rabbit IgG (H+L) | Jackson ImmunoResearch Laboratories, Inc. | RRID:AB_2492288 |
| **Bacterial and Virus Strains** |  |  |
| AAV9-CaMKIIa-eNpHR3.0-eYFP | Addgene | RRID:Addgene_26971 |
| AAV9-Ef1a-DIO-eNpHR3.0-eYFP | Addgene | RRID:Addgene_26966 |
| AAV9.EF1a.DIO.hChr2(H134R)-eYFP.WPRE.hGH | Addgene | RRID:Addgene_20298 |
| AAV9-hSyn-eGFP | Addgene | RRID:Addgene_50465 |
| **Chemicals** |  |  |
| Dil red | Thermo Fisher Scientific | Cat#D282 |
| **Experimental Models: Organisms/Strains** |  |  |
| Mouse: C57BL/6J | The Jackson Laboratory | RRID:IMSR_JAX:000664 |
| Mouse: B6.129S1-Htt<sup>tm1.1Mfc</sup>/190ChdJ | The Jackson Laboratory | RRID:IMSR_JAX:029928 |
| Mouse: B6J.129S6(FVB)-Slc17a6<sup>tm2(cre)Low</sup>/MwarJ | The Jackson Laboratory | RRID:IMSR_JAX:028863 |
| Mouse: B6.Cg-Pvalb<sup>tm1.1(cre)Aibs</sup>/J | The Jackson Laboratory | RRID:IMSR_JAX:012358 |
| **Software and Algorithms** |  |  |
| DeepLabCut | <sup>48</sup> | RRID:SCR_021391 |
| Fiji | https://imagej.net/software/fiji/ | RRID:SCR_003070 |
| Illustrator | Adobe | RRID:SCR_010279 |
| Mathematica | Wolfram Mathematica | RRID:SCR_014448 |
| MATLAB | MathWorks | RRID:SCR_001622 |
| NeuroExplorer | Nex Technologies | RRID:SCR_001818 |
| Plexon Offline Sorter software | Plexon, Inc. | RRID:SCR_000012 |
| Prism | GraphPad Software | RRID: SCR_002798 |
| R Project for Statistical Computing | The R Foundation | RRID:SCR_001905 |

## RESOURCE AVAILABILTY

### Lead Contact

Further information and requests for resources and reagents should be directed to and will be fulfilled by the lead contact, Mark Bevan.

### Materials Availability

All materials are available from the commercial vendors listed in the Key Resources Table.

### Data and Code Availability

Original datasets are available upon request to the lead contact.

## EXPERIMENTAL MODEL AND STUDY PARTICIPANT DETAILS

### Animals

Adult male and female heterozygous Q175 mice (B6.129S1-Htt^tm1.1Mfc^/190ChdiJ; RRID:IMSR_JAX:029928; The Jackson Laboratory, Bar Harbor, ME, USA) and their wild-type (WT; C57B6/J; RRID:IMSR_JAX:000664; The Jackson Laboratory) littermates were used in this study (Q175: 251.0, 225.0-264.5 days old, n = 5 male, 4 female; WT: 223.0, 211.3-232.8 days old, n = 6 male, 6 female). For optogenetic experiments, adult male and female heterozygous Vglut2-ires-cre knock-in (C57BL/6J) mice (B6J.129S6(FVB)-Slc17a6^tm2(cre)Low^/MwarJ; RRID:IMSR_JAX:028863; The Jackson Laboratory; 139, 134.0-187.5 days old; n = 1 male, 3 female) and PV-cre mice (B6.Cg-Pvalb^tm1.1(cre)Aibs^/J; RRID:IMSR_JAX:012358; The Jackson Laboratory; 145.5, 135.8-147.8 days old; n = 2 male, 2 female) were used. Data from male and female mice were overlapping and therefore pooled. Mice of the same sex were housed 1–5 per cage, maintained on a 14 h light/10 h dark cycle (conventional light cycle) or a 12 h dark/12 h light cycle (reverse light cycle) with food and water available ad libitum. Measurements from mice in the dark (active) and light (inactive) phases of their light cycle were overlapping and therefore pooled, except for kinematic measurements, which were only made in mice during the dark (active) phases of their light cycle. Mice were monitored regularly by animal care technicians, veterinarians, and research staff. All procedures were performed in compliance with the policies of the National Institutes of Health and approved by the Institutional Animal Care and Use Committee of Northwestern University.

## METHOD DETAILS

### Stereotaxic injection of viral vectors (dx.doi.org/10.17504/protocols.io.q26g7yr78gwz/v1)

Before and after surgical procedures, lab surfaces and equipment were disinfected with 5% Nolvasan Surgical Scrub (Zoetis, Inc., Kalamazoo, MI, USA). All surgical instruments were autoclaved, and sterility was maintained throughout surgery. Anesthesia was induced with vaporized 3-4% isoflurane (Smiths Medical ASD, Inc., Dublin, OH, USA) followed by an intraperitoneal injection of ketamine (100 mg/kg in 0.9% saline solution). The fur overlying the dorsal surface of the head was shaved using hair clippers (Wahl, Sterling, IL, USA). Ophthalmic ointment (Covetrus, Portland, ME, USA) was applied to prevent corneal drying. The mouse was transferred to a stereotaxic apparatus (Neurostar, Tübingen, Germany) and placed on a thermal heating pad (Physitemp, Clifton, NJ, USA) covered in a sterile drape. The animal’s head was secured in the stereotaxic frame with ear bars and anesthesia was maintained with 1-2% isoflurane. Depth of anesthesia was assessed by toe pinch throughout the procedure. The scalp was disinfected with 70% ethanol (Medline, Northfield, IL, USA) and Betadine (Dynarex Corporation, Orangeburg, NY, USA) and the skull was exposed with a single scalpel cut along the midline. AAVs diluted in sterile-filtered HEPES-buffered synthetic interstitial fluid (HBS SIF: 140 mm NaCl, 23 mm glucose, 15 mm HEPES, 3 mm KCl, 1.5 mm MgCl2, 1.6 mm CaCl_2_; pH 7.2 with NaOH; 300-310 mOsm/L) were then injected under stereotaxic guidance. eNpHR3.0-eYFP was virally expressed in the STN of WT mice through unilateral injection of AAV9-CaMKIIa-eNpHR3.0-eYFP (RRID:Addgene_26971; diluted to 2.4-4.8 x 10^12^ genome copies (GC)/mL; coordinates relative to bregma, AP: -1.90 mm; ML: 1.60 mm; DV: 4.60 mm; 0.3 μl). eNpHR3.0-eYFP was also virally expressed in the STN of vGluT2-Cre mice through unilateral injection of Cre-dependent AAV9-Ef1a-DIO-eNpHR3.0-eYFP (RRID:Addgene_26966; diluted to 1.9-4.4 x 10^12^ genome copies (GC)/mL; coordinates relative to bregma, AP: -1.90 mm; ML: 1.60 mm; DV: 4.60 mm; 0.3 μl). eNpHR3.0-eYFP or ChR2(H134R)-eYFP was virally expressed in parvalbumin-expressing STN neurons in PV-Cre mice through unilateral injection of Cre-dependent AAV9-Ef1a-DIO eNpHR3.0-eYFP (RRID:Addgene_26966; diluted to 1.9-4.4 x 10^12^ genome copies (GC)/mL; coordinates relative to bregma, AP: -1.90 mm; ML: 1.60 mm; DV: 4.60 mm; 0.3 μl) or AAV9.EF1a.DIO.hChR2(H134R)-eYFP.WPRE.hGH (RRID:Addgene_20298; diluted to 2.4 x 10^12^ genome copies (GC)/mL; coordinates relative to bregma, AP: −1.90 mm; ML: 1.60 mm; DV: 4.60 mm; 0.3 μl), respectively. Finally, eGFP was virally expressed in STN neurons through unilateral injection of AAV9-hSyn-eGFP (RRID:Addgene_50465; diluted to 2.0 x 10^12^ genome copies (GC)/mL; coordinates relative to bregma, AP: −1.90 mm; ML: 1.60 mm; DV: 4.60 mm; 0.3 μl). AAV injections were conducted over 10 min. The injectate was then allowed to diffuse for 10 minutes before the syringe was slowly retracted. The scalp was then sutured using nylon surgical sutures (6-0) (Henry Schein, Melville, NY, USA) and an analgesic, meloxicam (20 mg/kg in 0.9% saline solution) was administered by subcutaneous injection. After surgery mice were removed from the stereotaxic instrument and returned to their home cage. Cages were placed on an electric heating pad (Sunbeam Products, Boca Raton, FL, USA) until animals were ambulatory. The period between AAV injection and neuronal recording was between 2-4 weeks.

### Headplate implantation (dx.doi.org/10.17504/protocols.io.bp2l694z5lqe/v1)

Each mouse was prepared and anesthesia was induced, maintained, and assessed as for AAV injection. The skin overlying the cranium was removed using fine surgical scissors (Fine Science Tools, Foster City, CA, USA) such that the cranial suture points between bregma and lambda were exposed. Cotton tipped applicators were used to peel away the periosteum from the exposed cranium. To enhance the bonding of dental cement (Parkell, Inc., Edgewood, NY, USA) to the cranium, the surface of the skull was then gently scored with a handheld micromotor drill (Stoelting Co, Wood Dale, IL, USA). The micromotor drill was then used to drill a burr hole in the cranium overlying above the STN (coordinates relative to bregma, AP: −1.90 mm; ML: 1.60 mm). A thin layer of Kwik-Sil silicone sealant (World Precision Instruments, Sarasota, FL, USA) was applied to bregma and the craniotomy overlying the STN to prevent the stereotaxic reference point and craniotomy from being exposed to dental cement. Kwik-Sil was allowed 1-2 min to cure before proceeding. A separate craniotomy was made above the cerebellum (coordinates relative to bregma, AP: -6.75 mm; ML: 0.80 mm). The dura was then carefully removed using a bent injection needle and a peridural screw electrode (MS51960-1; McMaster-Carr, Elmhurst, IL, USA) was affixed in place. A custom-designed titanium headplate (25.4 mm length x 9.4 mm width x 0.8 mm thickness) with a 5.8 mm × 3.0 mm oval opening was used for head-fixation. The headplate was attached to a custom holder mounted to the stereotaxic frame. A thin layer of dental cement was then applied to the skull and base of the peridural screw electrode to cover the exposed skull and provide a foundation for affixing the headplate. Another layer of dental cement was then applied to affix the headplate, leaving only the Kwik-Sil enclosed craniotomy above the STN and screw electrode exposed. Dental cement was allowed to dry and harden for 10 min. Following headplate implantation, an analgesic was administered and mice were allowed to recover as for AAV injection.

### Treadmill habituation and behavior (dx.doi.org/10.17504/protocols.io.14egn2xrqg5d/v1)

Seventy-two hours or more after head plate implantation and one week prior to *in vivo* electrophysiological recording, mice were habituated to head-fixation on a cylindrical or linear treadmill. The cylindrical treadmill comprised a Styrofoam cylinder (16.5 cm in diameter x 15 cm wide) with a stainless-steel axle (1257K46; McMaster-Carr). Ball bearings (4262T11; McMaster-Carr) affixed to the ends of the axle were used to reduce friction. An axle-mounted optical encoder (E2; US Digital, Vancouver, WA, USA) was used to sample treadmill velocity at 1000 Hz. The axle of the cylindrical treadmill was supported via custom mounts affixed to a vibration isolation table (Ametek TMC, Peabody, MA, USA). The linear treadmill (SpeedBelt; Phenosys, Berlin, Germany) consisted of a fabric belt (60 cm length x 7.5 cm width) wrapped around two wheels on which ball bearings were mounted to reduce friction. An optical motion sensor was used to sample treadmill velocity at 1000 Hz.

Experiments were initially conducted during the light period of the light-dark cycle but were subsequently transitioned to the dark period when mice are more active. Mice were briefly placed in an anesthetic induction chamber (2-3% isoflurane) and then quickly transferred to the cylindrical or linear treadmill. Headplates were secured to custom-made clamps on each side of the head. Head-fixation posts and clamps were fabricated from commercially available components (Thorlabs, Newton, NJ, USA, and Luigs & Neumann, Ratingen, Germany). The final position of the headplate was typically 2.5-3.0 cm above the treadmill.

Digital movies were collected using 2 high-definition cameras (BFS-U3-04S2M-CS; Teledyne FLIR, Wilsonville, OR, USA) mounted to the front and side (contralateral to electrophysiological recording sites) of head-fixed mice. Frames were captured at 100 fps (720 × 540 pixels) using SpinView software (Spinnaker SDK; Teledyne FLIR). An infrared light source (CMVision, Houston, TX, USA) was used for illumination.

For experiments conducted using the cylindrical treadmill, habituation consisted of 3 sessions on consecutive days. The first session lasted for 30 min and subsequent sessions lasted for 60 min. On the first day of habituation mice attempted frequent postural adjustments and ambulated in both the forward and reverse directions. For experiments conducted using the linear treadmill, habituation consisted of five sessions on consecutive days. The first session lasted for 30 min and subsequent sessions lasted for 60 min. Over the course of habituation mice more consistently exhibited stereotyped and longer duration forward locomotion. After the conclusion of each habituation session, the animal’s headplate was detached from the headplate clamps and the mouse was returned to its home cage.

### In vivo electrophysiology and optogenetics (dx.doi.org/10.17504/protocols.io.rm7vzbn94vx1/v1)

The week following treadmill habituation, *in vivo* electrophysiological experiments were performed. Mice were head-fixed as described above. Using fine point forceps (Dumont #7 Forceps, Curved, Dumostar, 0.17 x 0.1 mm, 11.5 cm; Fine Science Tools) Kwik-Sil was carefully detached from the skull to expose both bregma and the craniotomy overlying the STN. Dura was carefully removed from the STN craniotomy using a bent syringe needle (25G). The craniotomy was then irrigated with HBS to prevent dehydration of exposed cortical tissue. Extracellular single-unit recordings were acquired using silicon probes and optrodes (A1x32-Poly3-10mm-50-177-A32 and A1x32-Poly3-10mm-50-177-OA32LP; NeuroNexus Technologies, Ann Arbor, MI; ASSY-77 Acute 64 channel H5 probe; Cambridge NeuroTech, Cambridge, UK) connected to a 64-channel Digital Lynx data acquisition system via a unity gain headstage (Neuralynx, Bozeman, MT, USA). Probes/optrodes were attached to a single-axis motorized micromanipulator (Scientifica, Uckfield, United Kingdom) that was mounted onto a 3-axis stereotaxic manual manipulator (David Kopf Instruments, Tujunga, CA, USA). The motorized micromanipulator was used to move probes/optrodes in the vertical axis and controlled using a PatchPad and LinLab software (Scientifica). The reference channel of the headstage was connected to the peridural screw electrode overlying the cerebellum by an insulated wire and signals were sampled at 40 kHz, with a gain of 14x. Online digital finite impulse response filters were applied. Single-unit activity was bandpass-filtered between 200 and 9000 Hz, and local field potential signals were bandpass-filtered between 0.1 and 400 Hz. A 633 nm direct diode laser (LuxX+ 633-100; Omicron-Laserage Laserprodukte, Rodgau, Germany) or a 473 nm direct diode laser (LuxX+ 473-100; Omicron-Laserage Laserprodukte) were used as light sources for optogenetic stimulation. An Axon Digidata 1440A (Molecular Devices, San Jose, CA, USA) and Clampex 11 software (Molecular Devices) were used to record treadmill velocity and synchronize video capture, optogenetic light delivery, and electrophysiological recording. eNpHR3.0-eYFP-expressing STN neurons were optogenetically inhibited through delivery of 633 nm light (<6 mW) for a duration of 5 s. Stimulation was repeated 4 times with each trial of stimulation separated by 2 min. ChR2(H134R)-eYFP-expressing STN neurons were optogenetically stimulated using 10 ms pulses of 473 nm light (<6 mW) delivered at 0.2 Hz for 250 sec. Laser intensity was calibrated as power at the tip of the optrode before implantation and verified at the conclusion of each experiment. In order to histologically verify recording sites, probe/optrode tracks were visualized postmortem by lightly dipping silicon probes in a lipophilic florescent dye (DiI; 20 mg/ml in 50% acetone/methanol; D282; Thermo Fisher Scientific, Waltham, MA, USA) prior to implantation or by immunohistochemical detection of the microglial marker, Iba1 (FUJIFILM Wako Pure Chemical Corporation, Richmond, VA, USA; RRID:AB_839504).

### Immunohistochemistry (dx.doi.org/10.17504/protocols.io.14egn2xrpg5d/v1) and confocal imaging (dx.doi.org/10.17504/protocols.io.rm7vzbn9rvx1/v1)

Following electrophysiological recording mice were given a lethal dose of anesthetic and then perfused transcardially with ∼5-10 ml of 0.01 M phosphate buffered saline (PBS) (pH 7.4; P3813; MilliporeSigma, Burlington, MA, USA) followed by 15-30 ml of 4% paraformaldehyde (PFA) in 0.1 M phosphate buffer (PB), pH 7.4. Each brain was then removed and postfixed overnight in 4% PFA (in 0.1 M PB, pH 7.4) before being washed in PBS, blocked, and sectioned in the coronal or sagittal plane at 70 µm with a vibratome (VT1000S; Leica Biosystems Inc., Buffalo Grove, IL, USA). Sections were then processed for the immunohistochemical detection of NeuN, an antigen expressed by neurons that is commonly used to delineate brain structures. First, sections were washed in PBS and incubated for 48-72 h at 4°C in anti-NeuN (1:500; Abcam, Cambridge, United Kingdom; RRID:AB_10711040) and, if applicable, anti-Iba1 (1:1000) in PBS with 0.3% Triton X-100 (MilliporeSigma) and 2% normal donkey serum (Jackson ImmunoResearch Laboratories, Inc., West Grove, PA, USA). Then, sections were washed in PBS before being incubated for 90 min at room temperature in Alexa Fluor 405, 488-, 594-, or 647-conjugated donkey anti-mouse or anti-rabbit IgG (1:250; Thermo Fisher Scientific; RRID:AB_2884884; Jackson ImmunoResearch Laboratories; RRID:AB_2313584; RRID:AB_2340621; RRID: AB_2492288, RRID: AB_2340846, RRID: AB_2340854, RRID: AB_2340863) in PBS with 0.3% Triton X-100 and 2% normal donkey serum. Finally, sections were washed in PBS and mounted on glass slides with ProLong Diamond Antifade Reagent (P36965; Thermo Fisher Scientific, Waltham, MA, USA). Mountant was allowed to cure for at least 24 h before storage at 4°C or imaging. In a subset of PV-cre mice, in which PV+ STN neurons expressed eNpHR3.0-eYFP, adjacent sections of the STN were processed for the immunohistochemical detection of PV, as described above (primary antibody: 1:1000 guinea pig anti-PV; Synaptic Systems; RRID:AB_2156476; secondary antibody: 1:250 AlexaFluor-594 donkey anti-guinea pig IgG; Jackson ImmunoResearch Laboratories; RRID:AB_2340474). DiI and immunofluorescent labeling were visualized using confocal laser scanning microscopy (A1, A1R or AX R; Nikon Instruments Inc., Melville, NY, USA). Whole-brain confocal images with DiI- or Iba1-labeled electrode tracks were plotted in the Allen Institute Common Coordinate Framework (CCF) using NeuroInfo (MBF Bioscience, Williston, VT, USA).

## QUANTIFICATION AND STATISTICAL ANALYSIS

### Treadmill behavior

Head-fixed mice spontaneously transitioned between periods of rest and locomotion. A treadmill velocity ≥ 0.25 cm/s for ≥ 200 ms was defined as locomotion. The start of a locomotor bout was defined as the first time point that treadmill velocity reached or exceeded 0.25 cm/s. The end of a locomotor bout was defined as the first time point that treadmill velocity fell below 0.25 cm/s for ≥ 500 ms. For electrophysiological comparisons, the minimum duration of a locomotor bout was defined as ≥ 1 s to prevent inclusion of short duration or fractionated running. Epochs during which locomotion occurred were divided into discrete periods defined as “rest” (1-2 s before locomotion onset), “pre-locomotion” (1 s before locomotion onset), “locomotion” (duration of the locomotor bout, as defined above), “post-locomotion” (1 s after locomotion), and “rest” (1-2 s after locomotion). To allow full sampling of the rest and peri-locomotor periods defined above, only locomotor bouts that were separated by > 3 s were analyzed. 30 s periods of continuous rest were used to analyze baseline neuronal activity. To compare locomotion parameters in WT and Q175 mice, 25 randomly selected bouts from each mouse were analyzed. Locomotor velocity was calculated by measuring the average treadmill velocity over each locomotor bout. Locomotor bout duration was determined by measuring the time from initiation to termination of locomotion, as defined above. The effects of optogenetic inhibition of the STN were studied in mice at rest or during coincident self-initiated locomotion. The impact of optogenetic inhibition on coincident locomotion was measured from the start of optogenetic inhibition to the end of the locomotor bout or the end of optogenetic inhibition, whichever occurred first.

DeepLabCut (version 2.2.2) was used to track the movement of the contralateral forepaw and hindpaw during locomotion. To create the training set, the hindpaw and forepaw were labeled in 20 frames from each of 48 videos (960 total frames) in 6 mice. 95% of the labeled data was used for training and 5% of labeled data was held out for testing. A ResNet-50-based neural network with an imgaug data loader for 500,000 training iterations was used for training. The test error was: 2.61 pixels, train: 3.71 pixels (image size was 720 × 540 pixels). We then used a p-cutoff of 0.9 to condition the X, Y coordinates for future analysis. The network was then used to analyze videos across similar experimental settings. The x-axis displacement of the hind paw was used to track the locomotor cycle. The valley and peak within each successive cycle were determined and used to classify the stance/propulsive phase (valley to peak) and swing phase (peak to valley). For outcome measures we compared the time, length, and velocity of the stance and swing phases during locomotion. To compare the kinematics of locomotion in Q175 and WT mice locomotor bouts that were sustained and equivalent in duration (3-5 seconds) were analyzed. There was no significant difference in the duration of the locomotor bouts used for this analysis (WT: 3.9, 3.5-4.5 s, n = 61 bouts; Q175: 3.7, 3.4-4.3 s, n = 70 bouts; values represent median and interquartile range). For optogenetic experiments, the impact of optogenetic inhibition on paw kinematics during coincident locomotion was analyzed by comparing equivalent numbers of step cycles in the pre-stimulus and optogenetic inhibition periods.

### In vivo electrophysiological analysis

Putative single-unit activity was discriminated with Plexon Offline Sorter software (Plexon, Inc., Dallas, Texas, USA; RRID:SCR_000012) using a combination of template matching, principal component analysis, and manual clustering. For classification of a single unit, sorting had to meet the following inclusion criteria: (1) PCA clusters were significantly different (p < 0.05); (2) J3-statistic ≥ 1; (3) Davies Bouldin test statistic ≤ 0.5. In addition, a threshold of < 0.5% of interspike intervals under 2 ms was required for classification as a putative single unit (% interspike interval within 2 ms; WT: 0.074, 0.0-0.200, n = 99; Q175: 0.0, 0.0-0.117, n = 103; values represent median and interquartile range). Electrophysiological data were visually inspected in NeuroExplorer (Nex Technologies; RRID:SCR_001818) and then exported to MATLAB (MathWorks, Natick, MA, USA; RRID:SCR_001622) and Mathematica (Wolfram Mathematica; RRID:SCR_014448). Recording sessions with stable unit isolation for ≥ 2 min were selected for analysis. Mean firing rates were calculated as the reciprocal of the mean interspike interval (1/mean interspike interval). For periods with ≤ 1 interspike interval (ISI), mean firing rates were calculated as the number of spikes divided by epoch length. The coefficient of variation (CV) of the interspike interval was used as a metric of regularity. If there were less than 3 spikes, the CV could not be calculated and was not reported. Histograms of spike rates (bin = 50 ms) were calculated for the entire recording period. Locomotion, peri-locomotion, and rest periods were defined as described above. Z-scores were determined relative to rest using:

bin z-score = (bin spike count –mean spike bin count at rest) / SD of bin counts at rest

For all locomotor bouts that met the criteria defined above, firing metrics were calculated and then averaged across bouts for each individual neuron. For correlation analysis, the velocity (v(t)) signal from the treadmill encoder was cross-correlated (c(k)) with the smoothed firing rate (r(t)) estimate of STN single units using:

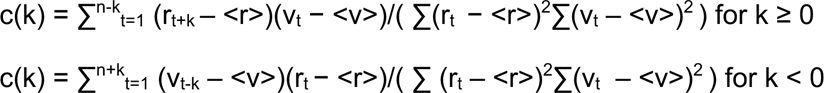

The firing rate, r(t), was estimated by smoothing the spike train, s(t), with a Gaussian function (SD = 0.04 s), G(t). Note that at zero lag (i.e. k = 0) this corresponds to Pearson’s correlation value. To determine the significance of the cross-correlation measurements, a spike-shuffled dataset was created using the spike times measured from recorded STN units. For each unit, a random draw from its ISI distribution was sampled without replacement using Mathematica’s function, RandomChoice[], and added to the first spike time to create the second spike time. Subsequently, another ISI was randomly drawn without replacement and added to the second spike time to create the third spike time, etc. This meant that the first spike time in the shuffled spike trains was always the same, but that subsequent addition of random draws from the ISI distribution created a random sequence of spike times. For each neuron, 20 shuffled spike trains were created, and each was cross-correlated with the smoothed velocity signal (Gaussian function (SD = 0.012 s)). Peak correlation values were measured and classified as significant if they fell outside of +/- 2 SDs from the average shuffled correlations. For functional classification of STN neurons, units were classified as “type 1,” “type 2,” or “uncorrelated”. Type 1 neurons had to exhibit both (1) a significant positive peak correlation and (2) a positive locomotion-associated z-score. Type 2 neurons had to exhibit both (1) a significant negative peak correlation and (2) a negative locomotion-associated z-score. If neurons failed to meet either of these conditions, they were categorized as uncorrelated.

To examine the relationship between neuronal firing and kinematics, phase histograms were generated in MATLAB. The instantaneous phase of each step cycle was calculated, and each spike was assigned to a phase of the step cycle from 0% to 100% (with 0% and 100% corresponding to the start and end of each cycle, respectively). Spikes phases were measured across cycles from each locomotor epoch and phase histograms were constructed using 10% bins. Spike times from each locomotor epoch were shuffled 1000 times and phase histograms were generated using the shuffled dataset. Neurons were considered phase-locked if their activity exceeded the shuffled mean by 2 SDs.

To determine whether neurons were responsive to optogenetic manipulation, peristimulus time histograms (PSTHs) were constructed from either 4 trials of eNpHR-EYFP stimulation or 50 trials of hChR2(H134R)-eYFP stimulation. VGLUT2- and PV-expressing STN neurons were considered directly responsive if their activity fell below 2 SDs of the pre-stimulus mean (5 s preceding stimulus onset) within 100 ms of eNpHR-EYFP stimulation (bin size 100 ms) and/or if spiking was silent throughout the 5 s stimulation period. PV+ STN neurons were considered directly responsive if their activity exceeded the pre-stimulus mean (100 ms preceding stimulus onset) by 2 SDs within 10 ms (bin size 1 ms) of hChR2(H134R)-eYFP stimulation.

### Experimental design

Data are reported as median and interquartile range. Data are represented graphically as violin (kernel density) plots and overlaid box plots, with the median (central line), interquartile range (box), and 10%-90% range (whiskers) denoted. The number and nature of observations for each parameter are specified throughout. To ensure that the proposed research was adequately powered, sample sizes were estimated using the formulae described by Noether ^94^ assuming 80% power (i.e., a 20% probability of a Type 2 error) and a two tailed α level of 0.05. For unpaired data (groups X and Y), and probabilities of X > Y (or X < Y) being 0.7, 0.8, and 0.9, the estimated sample sizes for each group are 33, 15, and 9, respectively. For paired data (where Xi and Xj are independent samples from X, reflecting effect size and sign) and the probabilities of Xi + Xj > 0 being 0.7, 0.8, and 0.9, the estimated sample sizes are 66, 30, and 17, respectively. Probabilities between 0.7 and 0.9 are representative of our historical and pilot data. To minimize assumptions concerning the distribution of data, nonparametric, two-tailed statistical comparisons were made using the Mann–Whitney U (MWU) and Wilcoxon signed-rank (WSR) tests for unpaired and paired comparisons, respectively. In addition, Fisher’s exact test was used for contingency analyses. P < 0.05 was considered significant. Where appropriate, p values were adjusted for multiple comparisons using the Holm–Bonferroni method. Plots and statistical comparisons were generated in Prism (GraphPad Software, San Diego, CA, USA; RRID:SCR_002798) and R (https://www.r-project.org/; RRID:SCR_001905).

## SUPPLEMENTAL INFORMATION

**Figure S1.**
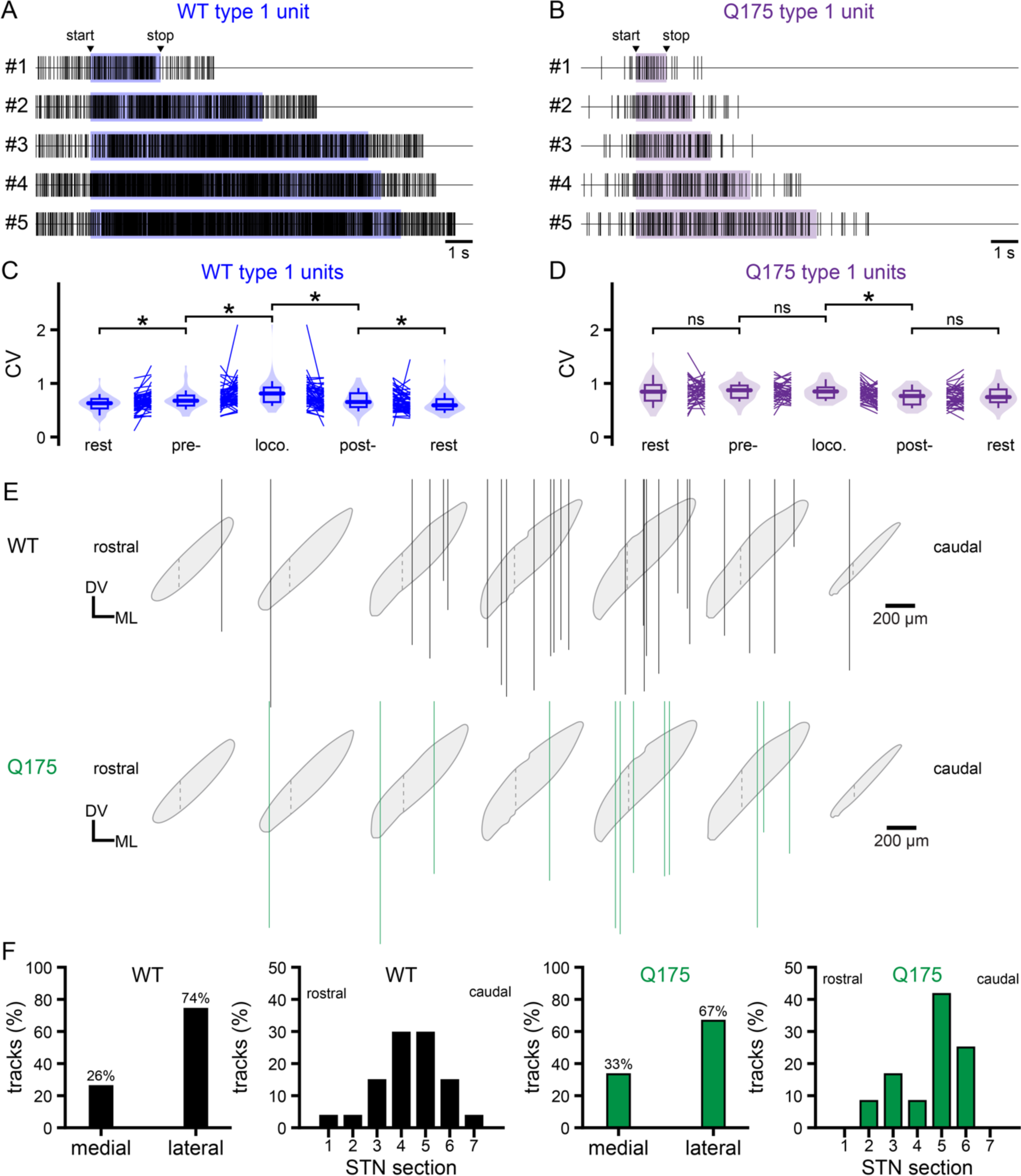
Related to Figure 3. Consistency, precision, and sampling of movement-related STN activity. (A-B) Representative examples of an individual type 1 unit’s activity in a WT (A) or Q175 (B) mouse across multiple locomotor bouts. (C-D) Population data. CV of type 1 unit activity in WT (C) and Q175 (D) mice during rest, pre-locomotor, locomotor, and post-locomotor periods. (E-F) Population data. Spatial location of electrode/optrode recording tracks that sampled movement-related STN unit activity in WT and Q175 mice. The boundary between the medial third and lateral two-thirds of the STN is denoted by a dashed line. *, p<0.05; ns, not significant.

**Figure S2.**
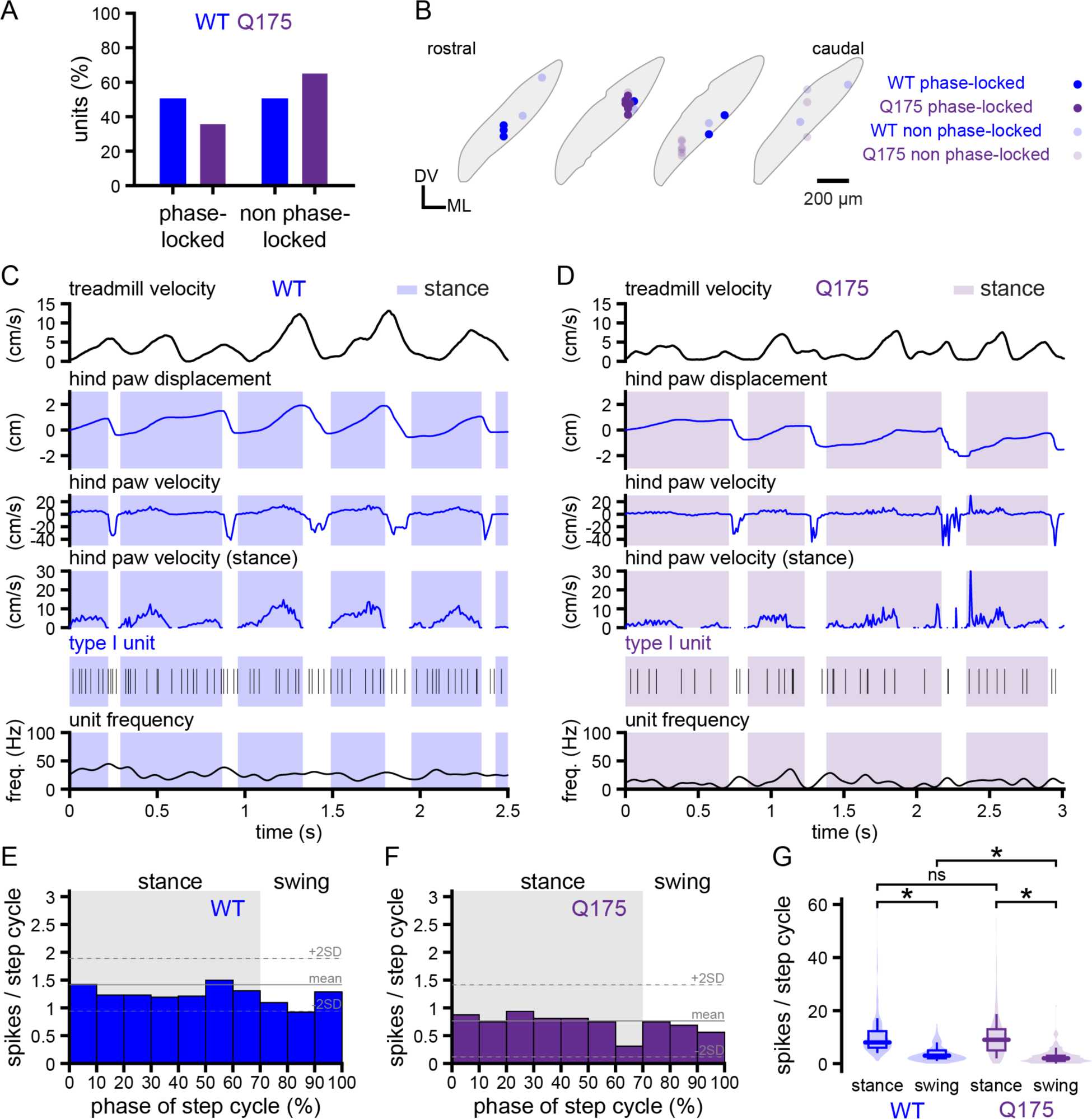
Related to Fig. 4. A subset of type 1 units exhibit activity that is unrelated to the phase of the locomotor cycle. (A-B) The distribution of type 1 neurons whose activity was or was not phase-locked to the locomotor cycle were distributed similarly in WT and Q175 mice. (C-F) As evinced by their associated phase histograms, a subset of type 1 STN neurons in WT and Q175 mice exhibited firing that was not related to the phase of locomotor cycle (C-F, representative examples). (G) During the swing phase, the number of spikes per cycle was lower in Q175 versus WT mice. *, p < 0.05; ns, not significant.

**Figure S3.**
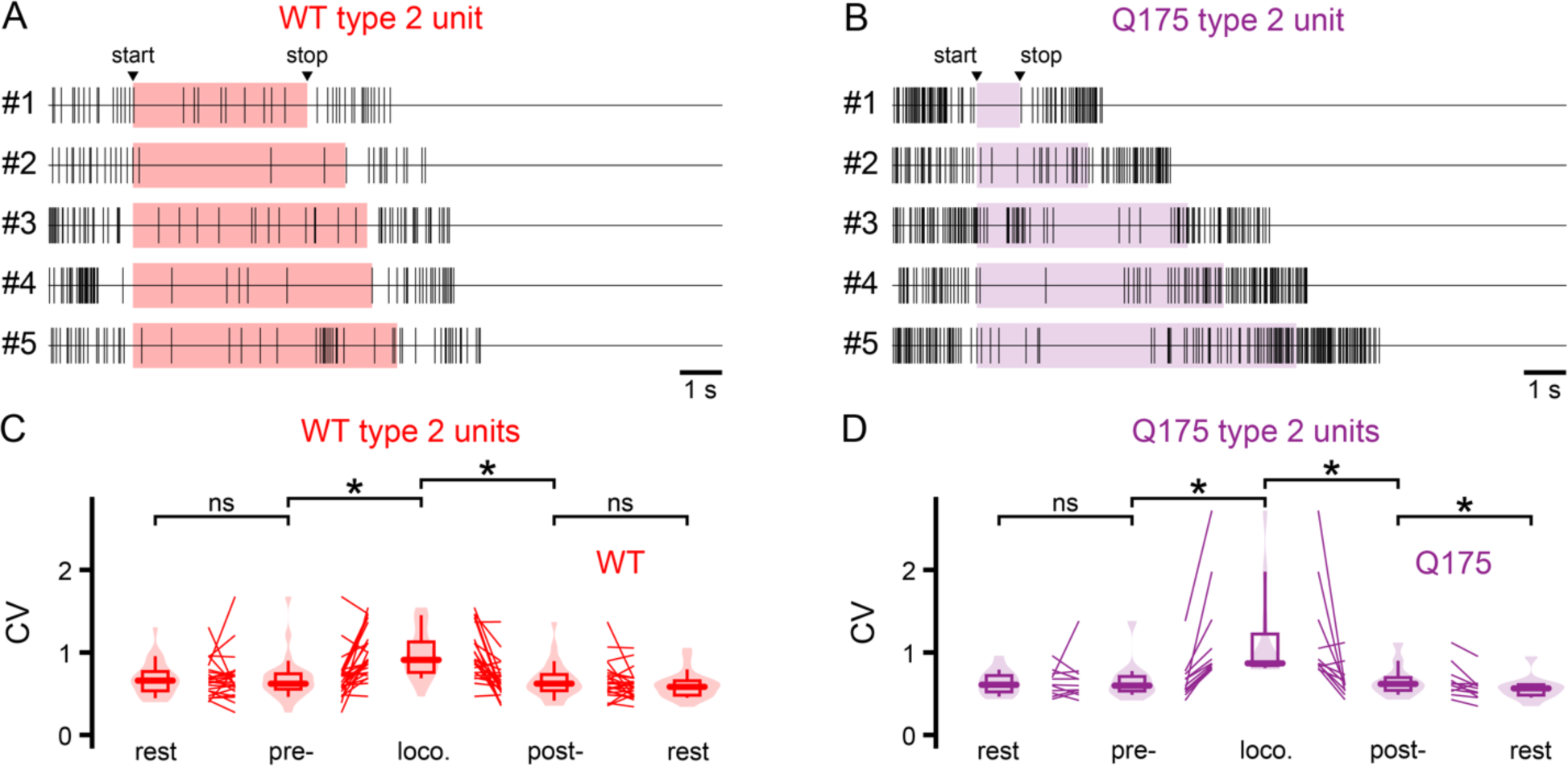
Related to Figure 5. Consistency and precision of movement-related type 2 STN activity. (A-B) Representative examples of an individual type 2 unit’s activity in a WT (A) or Q175 (B) mouse across multiple locomotor bouts. (C-D) Population data. CV of type 2 unit activity in WT (C) and Q175 (D) mice during rest, pre-locomotor, locomotor, and post-locomotor periods. *, p<0.05; ns, not significant.

**Figure S4.**
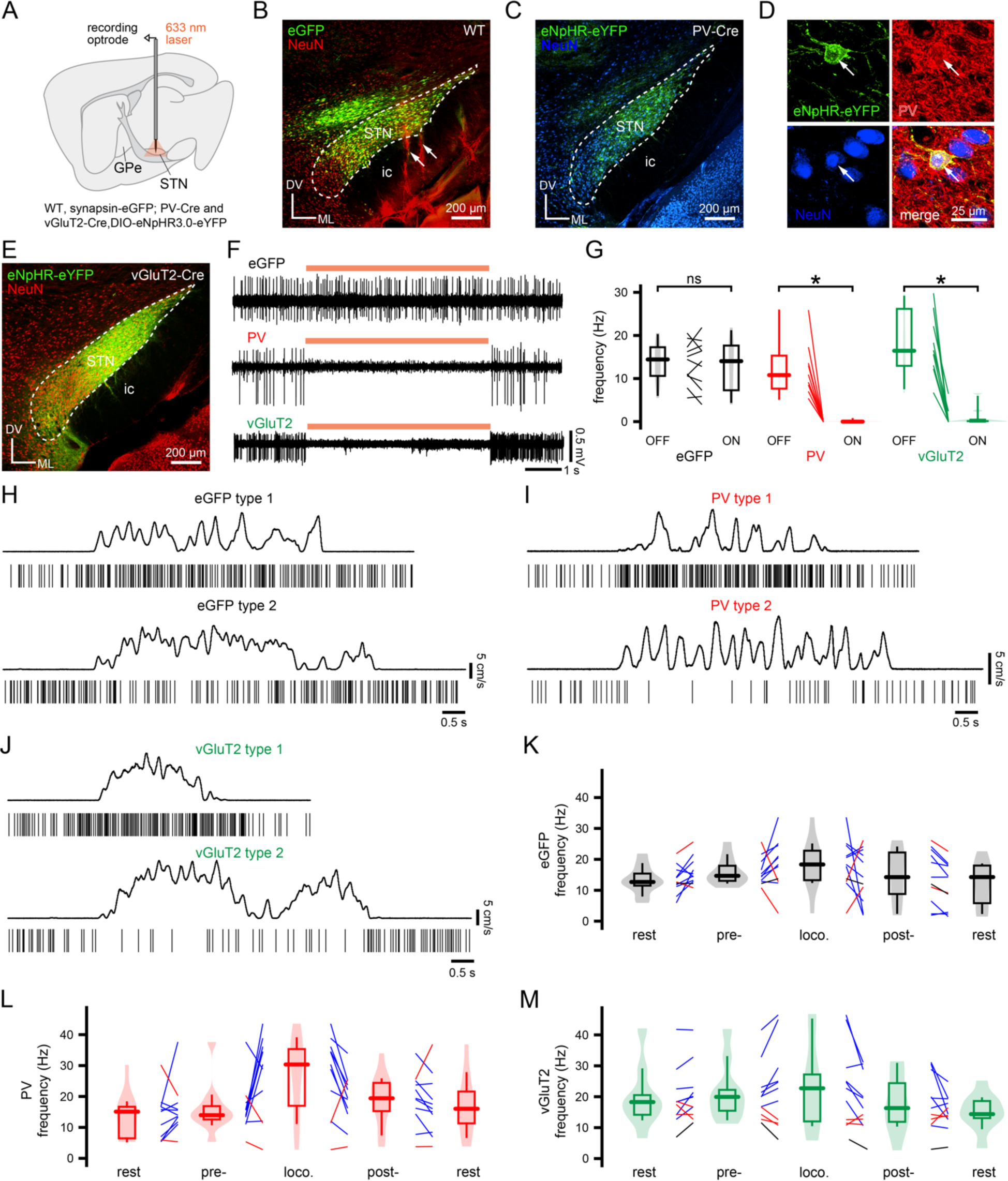
PV+ and vGluT2+ STN neurons exhibit heterogeneous locomotion encoding properties. (A-G) Experimental approach (A). eGFP (green) was expressed in the STN of WT control mice through intracerebral injection of AAVs carrying a synapsin-dependent construct (B, confocal micrograph of eGFP expression in the STN; electrode recording tracks (white arrows) are labeled with DiI red; ic, internal capsule; NeuN counterstain, red; DV, dorsoventral axis; ML, mediolateral axis). eNpHR3.0-eYFP (green) was expressed in PV+ STN neurons in PV-Cre mice through injection of AAVs carrying a Cre-dependent expression construct (C, eNpHR3.0-eYFP expression in PV+ STN neurons; NeuN counterstain, blue; D, eNpHR3.0-eYFP expression in an immunohistochemically identified PV+ STN neuron (red); NeuN counterstain, blue). eNpHR3.0-eYFP (green) was expressed in vGluT2-expressing STN neurons in vGluT2-Cre mice through injection of AAVs carrying a Cre-dependent expression construct (E, eNpHR3.0-eYFP expression in vGluT2+ STN neurons; NeuN counterstain, red). Delivery of 633 nm light had no effect on STN unit activity in eGFP-expressing control mice (F, representative example; G, population data). In eNpHR3.0-expressing PV+ and vGluT2+ STN neurons, activation of the inhibitory opsin through 633 nm light delivery rapidly and powerfully suppressed unit activity (F, representative examples; G, population data). (H-M) PV+ and vGluT2+ STN neurons exhibited type 1 or type 2 encoding properties in approximately the same proportion as STN neurons in eGFP-expressing control mice (H-J, representative examples; upper panels, treadmill velocity; lower panels, examples of type 1 and 2 unit activity before, during, and following treadmill locomotion; K-M, population data; blue, type 1 units; red, type 2 units; black, uncorrelated units). (A-B) STN neurons in eGFP-expressing control mice were unresponsive to delivery of 633 nm light for 5 seconds (A, population peristimulus time histogram. Orange bar denotes 633 nm light delivery for 5 seconds) and exhibited diverse locomotion-encoding properties (B, Population data. Z-scores of locomotion-related unit activity in control mice; blue, type 1; red, type 2; black, uncorrelated). (C-F) PV+ STN neurons were optotagged through activation of eNpHR3.0-eYFP (C, population peristimulus time histogram) or ChR2(H134R)-eYFP (D, schematic of experimental approach; E, representative peristimulus time histogram. Blue bar denotes 473 nm light delivery for 5 ms) and exhibited diverse locomotion-encoding properties (F, population data. Z-scores of locomotion-related PV+ unit activity; blue, type 1; red, type 2). (G-H) vGluT2+ STN neurons were optotagged through activation of eNpHR3.0-eYFP (G, population peristimulus time histogram. Orange bar denotes 633 nm light delivery for 5 seconds) and exhibited diverse locomotion-encoding properties (H, population data. Z-scores of locomotion-related vGluT2+ unit activity; blue, type 1; red, type 2; black, uncorrelated). (I-M) The proportions (I, population data; 1, type 1; 2, type 2; U, uncorrelated) and spatial distributions (J, population data) of control and optotagged PV+ and vGluT2+ neurons exhibiting type 1 or type 2 neuronal activity were similar. The baseline firing frequencies (K) and locomotion-encoding properties (L, locomotion-associated firing frequencies; M, locomotion-associated z-scores) of control and optotagged PV+ and vGluT2+ neurons were also similar. *, p<0.05; ns, not significant.

**Figure S5.**
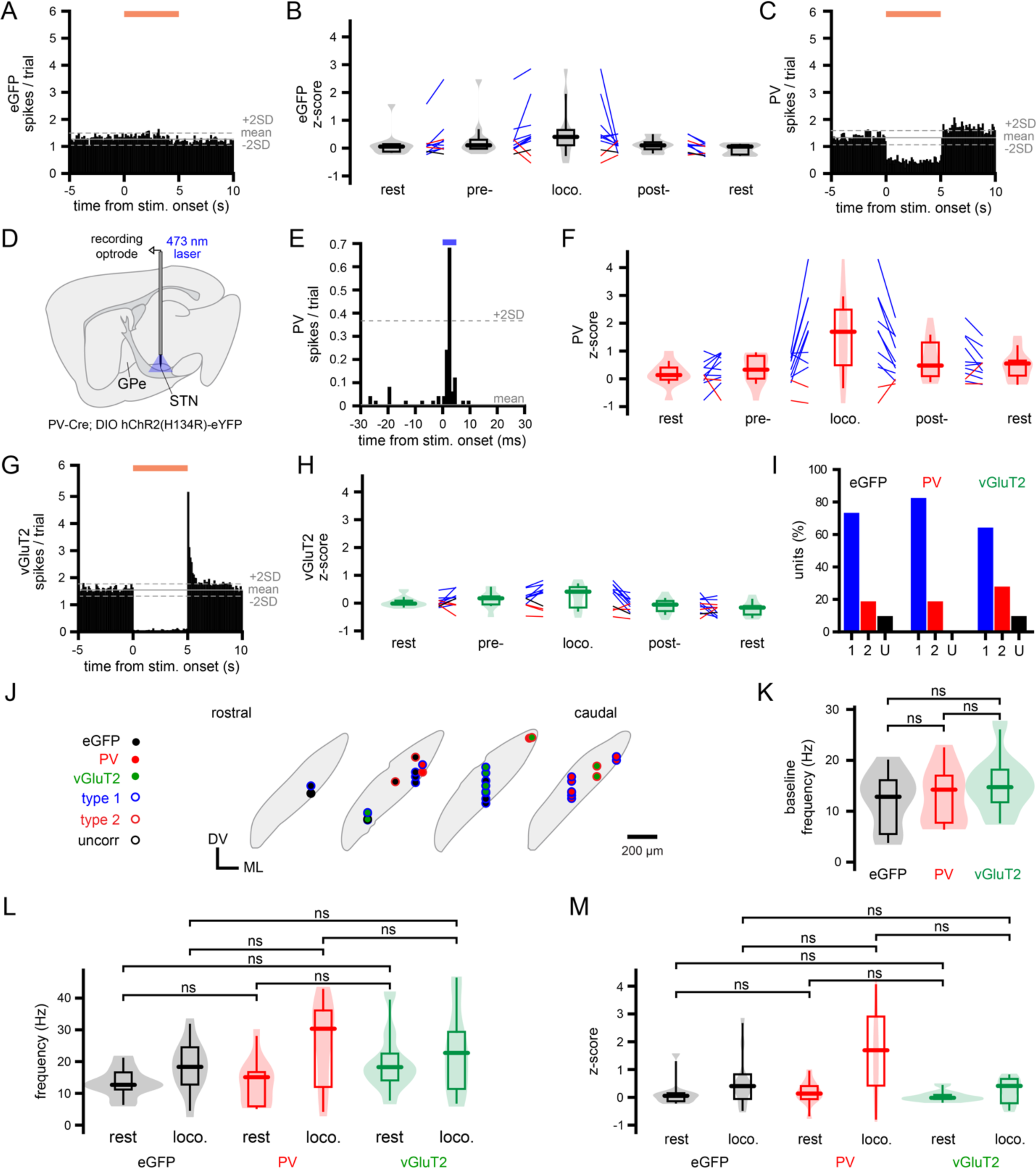
Locomotion-encoding properties of STN subtypes. (A) Experimental approach (A). eNpHR3.0-eYFP was expressed in the STN of WT control mice through intracerebral injection of AAVs carrying a CaMKII-dependent expression construct. (B) Confocal micrograph of eNpHR3.0-eYFP expression in the STN (electrode recording tracks (white arrows) are labeled with DiI red; ic, internal capsule; NeuN counterstain, red; DV, dorsoventral axis; ML, mediolateral axis). (C-D) In eNpHR3.0-eYFP-expressing STN neurons, activation of the inhibitory opsin through delivery of 633 nm light (denoted by orange bar) rapidly and powerfully suppressed unit activity (C, representative trace; D, associated peristimulus time histogram). (E) Population data. Activation of eNpHR3.0-eYFP expressed in WT mice using a CAMKII-promoter-dependent construct or in vGluT2-Cre mice using a Cre-dependent construct similarly suppressed STN activity. (F) Population data. eNpHR3.0-eYFP activation similarly suppressed the activity of type 1, type 2, and uncorrelated STN units. (G) Population data. eNpHR3.0-eYFP-mediated optogenetic inhibition of the lateral and middle thirds of the STN truncated locomotor bout duration relative to light delivery alone in eGFP-expressing mice or eNpHR3.0-eYFP-mediated optogenetic inhibition of the middle third of the STN. (H) Population data. Activation of eNpHR3.0-eYFP expressed in WT mice using a CAMKII-promoter-dependent construct or in vGluT2-Cre mice using a Cre-dependent construct similarly truncated locomotor bout duration. (I) Population data. The durations of locomotor bouts concomitant with eNpHR3.0-eYFP-mediated optogenetic inhibition of the STN during the first 50% of trials and the last 50% of trials of light delivery were not significantly different. *, p<0.05; ns, not significant; n/a, not applicable.

**Figure S6.**
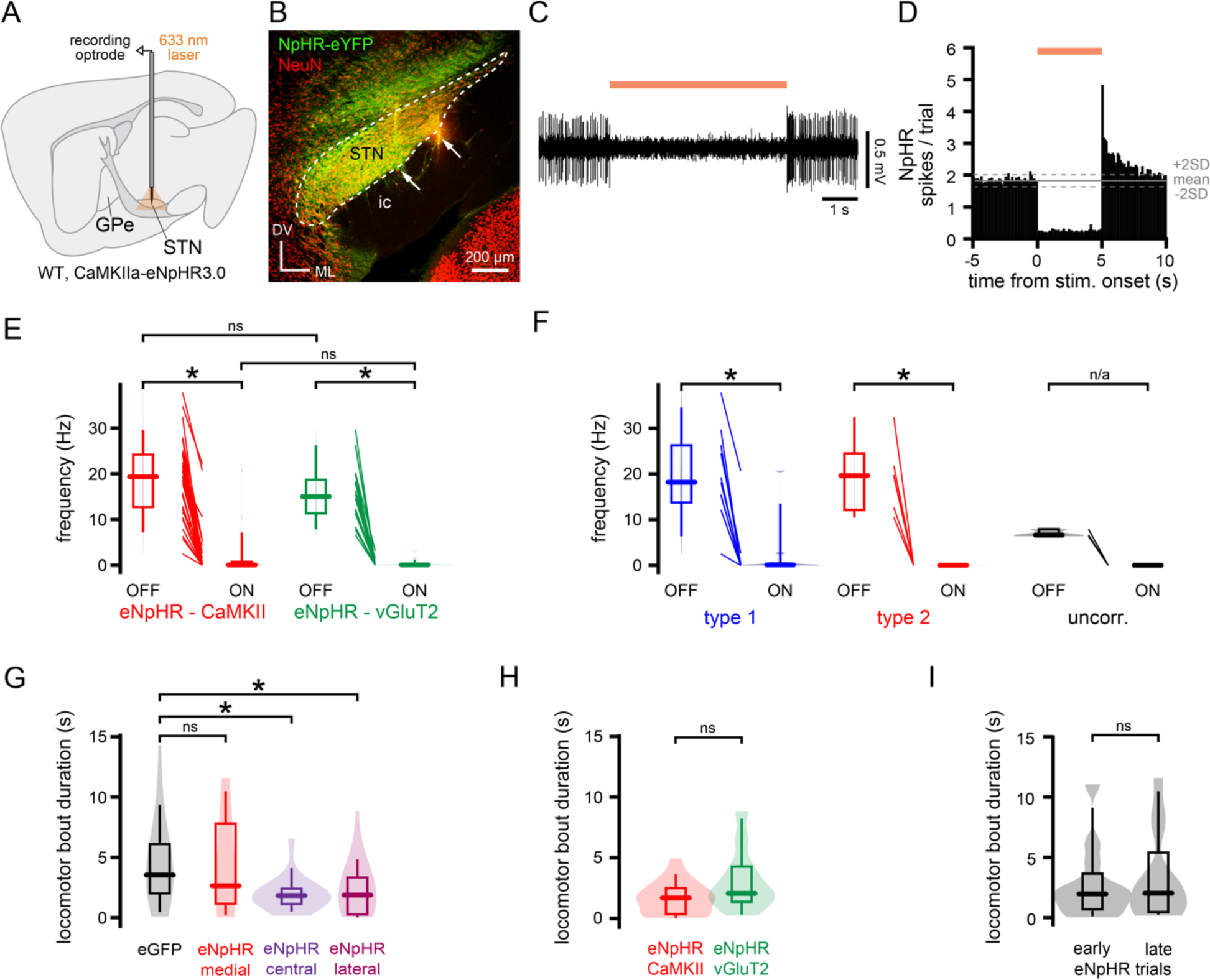
Related to Figure 7. eNpHR3.0-eYFP-mediated optogenetic inhibition of the STN.

**Table S1.**
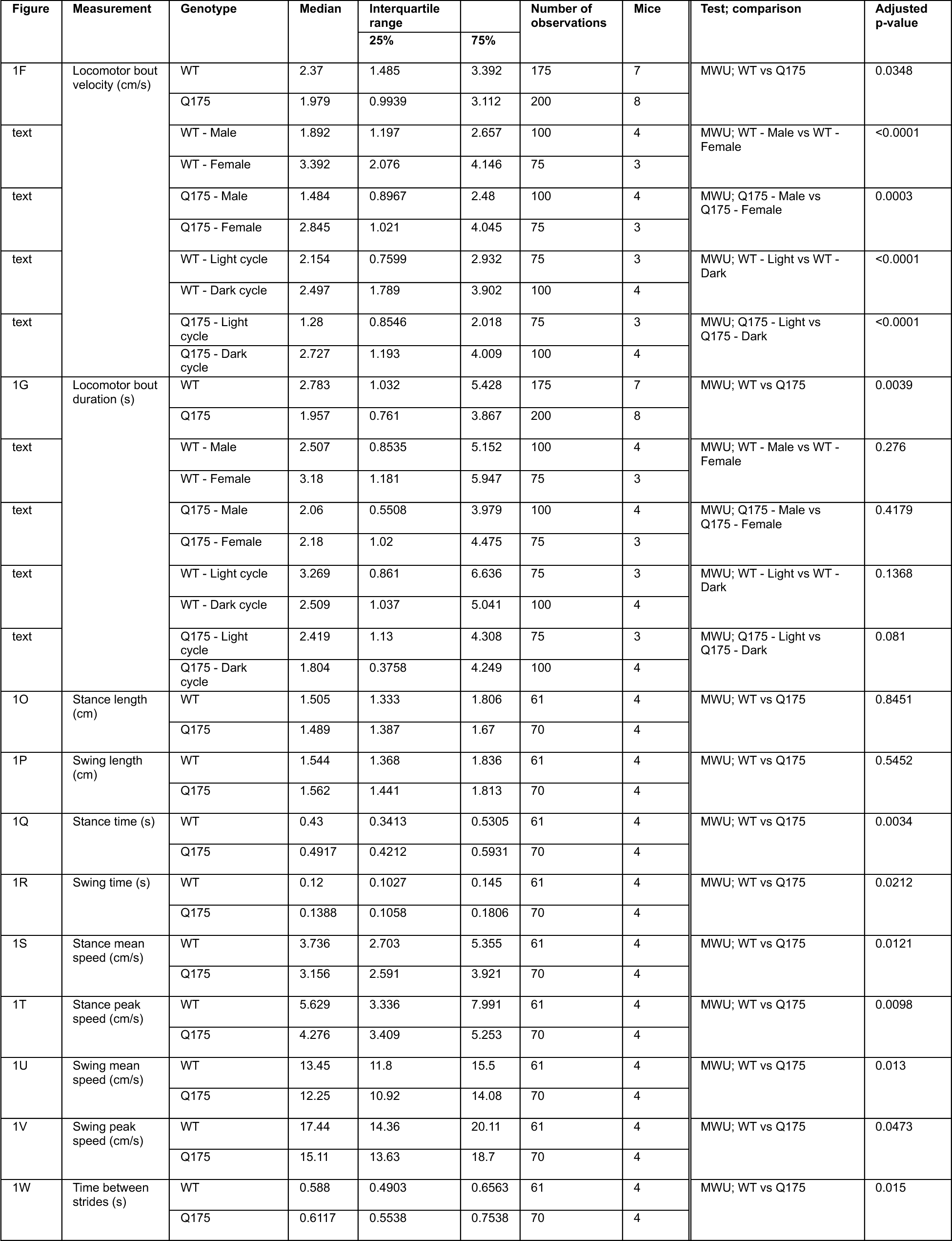
Self-initiated locomotion is dysregulated in Q175 HD mice. Related to Figure 1.

| Figure | Measurement | Genotype | Median | Interquartile range |  | Number of observations | Mice | Test; comparison | Adjusted p-value |
| --- | --- | --- | --- | --- | --- | --- | --- | --- | --- |
|  |  |  |  | 25% | 75% |  |  |  |  |
| 1F | Locomotor bout velocity (cm/s) | WT | 2.37 | 1.485 | 3.392 | 175 | 7 | MWU; WT vs Q175 | 0.0348 |
|  |  | Q175 | 1.979 | 0.9939 | 3.112 | 200 | 8 |  |  |
| text |  | WT - Male | 1.892 | 1.197 | 2.657 | 100 | 4 | MWU; WT - Male vs WT - Female | <0.0001 |
|  |  | WT - Female | 3.392 | 2.076 | 4.146 | 75 | 3 |  |  |
| text |  | Q175 - Male | 1.484 | 0.8967 | 2.48 | 100 | 4 | MWU; Q175 - Male vs Q175 - Female | 0.0003 |
|  |  | Q175 - Female | 2.845 | 1.021 | 4.045 | 75 | 3 |  |  |
| text |  | WT - Light cycle | 2.154 | 0.7599 | 2.932 | 75 | 3 | MWU; WT - Light vs WT - Dark | <0.0001 |
|  |  | WT - Dark cycle | 2.497 | 1.789 | 3.902 | 100 | 4 |  |  |
| text |  | Q175 - Light cycle | 1.28 | 0.8546 | 2.018 | 75 | 3 | MWU; Q175 - Light vs Q175 - Dark | <0.0001 |
|  |  | Q175 - Dark cycle | 2.727 | 1.193 | 4.009 | 100 | 4 |  |  |
| 1G | Locomotor bout duration (s) | WT | 2.783 | 1.032 | 5.428 | 175 | 7 | MWU; WT vs Q175 | 0.0039 |
|  |  | Q175 | 1.957 | 0.761 | 3.867 | 200 | 8 |  |  |
| text |  | WT - Male | 2.507 | 0.8535 | 5.152 | 100 | 4 | MWU; WT - Male vs WT - Female | 0.276 |
|  |  | WT - Female | 3.18 | 1.181 | 5.947 | 75 | 3 |  |  |
| text |  | Q175 - Male | 2.06 | 0.5508 | 3.979 | 100 | 4 | MWU; Q175 - Male vs Q175 - Female | 0.4179 |
|  |  | Q175 - Female | 2.18 | 1.02 | 4.475 | 75 | 3 |  |  |
| text |  | WT - Light cycle | 3.269 | 0.861 | 6.636 | 75 | 3 | MWU; WT - Light vs WT - Dark | 0.1368 |
|  |  | WT - Dark cycle | 2.509 | 1.037 | 5.041 | 100 | 4 |  |  |
| text |  | Q175 - Light cycle | 2.419 | 1.13 | 4.308 | 75 | 3 | MWU; Q175 - Light vs Q175 - Dark | 0.081 |
|  |  | Q175 - Dark cycle | 1.804 | 0.3758 | 4.249 | 100 | 4 |  |  |
| 1O | Stance length (cm) | WT | 1.505 | 1.333 | 1.806 | 61 | 4 | MWU; WT vs Q175 | 0.8451 |
|  |  | Q175 | 1.489 | 1.387 | 1.67 | 70 | 4 |  |  |
| 1P | Swing length (cm) | WT | 1.544 | 1.368 | 1.836 | 61 | 4 | MWU; WT vs Q175 | 0.5452 |
|  |  | Q175 | 1.562 | 1.441 | 1.813 | 70 | 4 |  |  |
| 1Q | Stance time (s) | WT | 0.43 | 0.3413 | 0.5305 | 61 | 4 | MWU; WT vs Q175 | 0.0034 |
|  |  | Q175 | 0.4917 | 0.4212 | 0.5931 | 70 | 4 |  |  |
| 1R | Swing time (s) | WT | 0.12 | 0.1027 | 0.145 | 61 | 4 | MWU; WT vs Q175 | 0.0212 |
|  |  | Q175 | 0.1388 | 0.1058 | 0.1806 | 70 | 4 |  |  |
| 1S | Stance mean speed (cm/s) | WT | 3.736 | 2.703 | 5.355 | 61 | 4 | MWU; WT vs Q175 | 0.0121 |
|  |  | Q175 | 3.156 | 2.591 | 3.921 | 70 | 4 |  |  |
| 1T | Stance peak speed (cm/s) | WT | 5.629 | 3.336 | 7.991 | 61 | 4 | MWU; WT vs Q175 | 0.0098 |
|  |  | Q175 | 4.276 | 3.409 | 5.253 | 70 | 4 |  |  |
| 1U | Swing mean speed (cm/s) | WT | 13.45 | 11.8 | 15.5 | 61 | 4 | MWU; WT vs Q175 | 0.013 |
|  |  | Q175 | 12.25 | 10.92 | 14.08 | 70 | 4 |  |  |
| 1V | Swing peak speed (cm/s) | WT | 17.44 | 14.36 | 20.11 | 61 | 4 | MWU; WT vs Q175 | 0.0473 |
|  |  | Q175 | 15.11 | 13.63 | 18.7 | 70 | 4 |  |  |
| 1W | Time between strides (s) | WT | 0.588 | 0.4903 | 0.6563 | 61 | 4 | MWU; WT vs Q175 | 0.015 |
|  |  | Q175 | 0.6117 | 0.5538 | 0.7538 | 70 | 4 |  |  |

**Table S2.** The frequency and precision of STN unit activity are lower in Q175 HD versus WT mice at rest. Related to Figure 2.

| Figure | Measurement | Genotype | Median | Interquartile range |  | Number of observations | Mice | Test; comparison | Adjusted p-value |
| --- | --- | --- | --- | --- | --- | --- | --- | --- | --- |
|  |  |  |  | 25% | 75% |  |  |  |  |
| 2B | Frequency (Hz) | WT | 14.99 | 10.61 | 20.71 | 99 | 19 | MWU; WT vs Q175 | <0.0001 |
|  |  | Q175 | 8.567 | 1.906 | 18.38 | 103 | 8 |  |  |
| text |  | WT - Male | 14.82 | 11.09 | 21.93 | 48 | 9 | MWU; WT - Male vs WT - Female | 1.306 |
|  |  | WT - Female | 15.18 | 8.801 | 19.08 | 51 | 10 |  |  |
| text |  | Q175 - Male | 8.567 | 2.134 | 20.05 | 43 | 4 | MWU; Q175 - Male vs Q175 - Female | 0.7926 |
|  |  | Q175 - Female | 8.888 | 1.612 | 18.27 | 60 | 4 |  |  |
| text |  | WT - Light cycle | 15.67 | 11.35 | 23.87 | 46 | 4 | MWU; WT - Light vs WT - Dark | 0.1778 |
|  |  | WT - Dark cycle | 14.72 | 8.81 | 18.23 | 53 | 15 |  |  |
| text |  | Q175 - Light cycle | 8.051 | 1.703 | 17.37 | 74 | 5 | MWU; Q175 - Light vs Q175 - Dark | 0.4327 |
|  |  | Q175 - Dark cycle | 9.846 | 2.091 | 20.39 | 29 | 3 |  |  |
| 2D | CV | WT | 0.7831 | 0.6481 | 0.9925 | 99 | 19 | MWU; WT vs Q175 | <0.0001 |
|  |  | Q175 | 1.082 | 0.8251 | 1.371 | 102 | 8 |  |  |
| text |  | WT - Male | 0.7767 | 0.6088 | 0.9451 | 48 | 9 | MWU; WT - Male vs WT - Female | 0.4512 |
|  |  | WT - Female | 0.8736 | 0.6677 | 1.031 | 51 | 10 |  |  |
| text |  | Q175 - Male | 1.122 | 0.8059 | 1.455 | 43 | 4 | MWU; Q175 - Male vs Q175 - Female | 0.3109 |
|  |  | Q175 - Female | 1.021 | 0.8466 | 1.284 | 59 | 4 |  |  |
| text |  | WT - Light cycle | 0.7647 | 0.5855 | 0.9354 | 46 | 4 | MWU; WT - Light vs WT - Dark | 0.051 |
|  |  | WT - Dark cycle | 0.8785 | 0.699 | 1.019 | 53 | 15 |  |  |
| text |  | Q175 - Light cycle | 1.073 | 0.7554 | 1.377 | 73 | 5 | MWU; Q175 - Light vs Q175 - Dark | 0.3675 |
|  |  | Q175 - Dark cycle | 1.096 | 0.9271 | 1.397 | 29 | 3 |  |  |
| text | CV (STN freq. > 5 Hz) | WT | 0.7796 | 0.6287 | 0.9741 | 96 | 19 | MWU; WT vs Q175 | 0.0035 |
|  |  | Q175 | 0.9434 | 0.7175 | 1.117 | 61 | 8 |  |  |

**Table S3.** Type 1 STN neurons in WT and Q175 mice exhibit locomotion-associated increases in firing. Related to Figures 3 and S1.

| Figure | Measurement | Genotype | Behavioral state | Median | Interquartile range |  | Number of observations | Mice | Test; comparison | Adjusted p-value |
| --- | --- | --- | --- | --- | --- | --- | --- | --- | --- | --- |
|  |  |  |  |  | 25% | 75% |  |  |  |  |
| 3F | Frequency (Hz) | WT | rest | 17.5 | 11.01 | 23.77 | 61 | 15 | WSR; rest vs pre-loco. | <0.0001 |
|  |  |  | pre-loco. | 21.12 | 14.8 | 29.98 |  |  | WSR; pre-loco. vs locomotion | <0.0001 |
|  |  |  | locomotion | 29.53 | 22.44 | 39.32 |  |  | WSR; locomotion vs post-loco. | <0.0001 |
|  |  |  | post-loco. | 20.6 | 14.28 | 27.95 |  |  | WSR; post-loco. vs rest | <0.0001 |
|  |  |  | rest | 17.86 | 11.68 | 24.55 |  |  |  |  |
| 3G | z-score | WT | rest | 0.07891 | -0.1535 | 0.1795 | 61 | 15 | WSR; rest vs pre-loco. | <0.0001 |
|  |  |  | pre-loco. | 0.274 | 0.05535 | 0.7153 |  |  | WSR; pre-loco. vs locomotion | <0.0001 |
|  |  |  | locomotion | 0.6741 | 0.4543 | 1.706 |  |  | WSR; locomotion vs post-loco. | <0.0001 |
|  |  |  | post-loco. | 0.1979 | -0.0552 | 0.5362 |  |  | WSR; post-loco. vs rest | <0.0001 |
|  |  |  | rest | 0.0711 | -0.1871 | 0.29 |  |  |  |  |
| 3M | Frequency (Hz) | Q175 | rest | 8.948 | 3.612 | 15.72 | 52 | 7 | WSR; rest vs pre-loco. | <0.0001 |
|  |  |  | pre-loco. | 16.71 | 8.799 | 27.44 |  |  | WSR; pre-loco. vs locomotion | 0.0201 |
|  |  |  | locomotion | 18.34 | 11.95 | 28.78 |  |  | WSR; locomotion vs post-loco. | <0.0001 |
|  |  |  | post-loco. | 9.299 | 6.694 | 20.66 |  |  | WSR; post-loco. vs rest | <0.0001 |
|  |  |  | rest | 7.619 | 2.473 | 17.02 |  |  |  |  |
| 3N | z-score | Q175 | rest | -0.03419 | -0.1409 | 0.08412 | 52 | 7 | WSR; rest vs pre-loco. | <0.0001 |
|  |  |  | pre-loco. | 0.4079 | 0.1414 | 0.7582 |  |  | WSR; pre-loco. vs locomotion | <0.0001 |
|  |  |  | locomotion | 0.7079 | 0.3415 | 1.368 |  |  | WSR; locomotion vs post-loco. | <0.0001 |
|  |  |  | post-loco. | 0.1646 | -0.0214 | 0.4499 |  |  | WSR; post-loco. vs rest | <0.0001 |
|  |  |  | rest | -0.0409 | -0.1933 | 0.1229 |  |  |  |  |
| S1C | CV | WT | rest | 0.6311 | 0.5302 | 0.7171 | 58 | 15 | WSR; rest vs pre-loco. | 0.0009 |
|  |  |  | pre-loco. | 0.677 | 0.5851 | 0.7831 |  |  | WSR; pre-loco. vs locomotion | 0.0004 |
|  |  |  | locomotion | 0.8116 | 0.6466 | 0.9289 |  |  | WSR; locomotion vs post-loco. | <0.0001 |
|  |  |  | post-loco. | 0.6535 | 0.5515 | 0.8236 |  |  | WSR; post-loco. vs rest | 0.0004 |
|  |  |  | rest | 0.5917 | 0.5133 | 0.7101 |  |  |  |  |
| S1D | CV | Q175 | rest | 0.8459 | 0.6688 | 0.9883 | 42 | 7 | WSR; rest vs pre-loco. | 0.9605 |
|  |  |  | pre-loco. | 0.8729 | 0.7279 | 0.9652 |  |  | WSR; pre-loco. vs locomotion | 1.7574 |
|  |  |  | locomotion | 0.8502 | 0.7293 | 0.9356 |  |  | WSR; locomotion vs post-loco. | 0.0072 |
|  |  |  | post-loco. | 0.7655 | 0.5996 | 0.8657 |  |  | WSR; post-loco. vs rest | 1.7574 |
|  |  |  | rest | 0.7452 | 0.6413 | 0.892 |  |  |  |  |

**Table S4.**
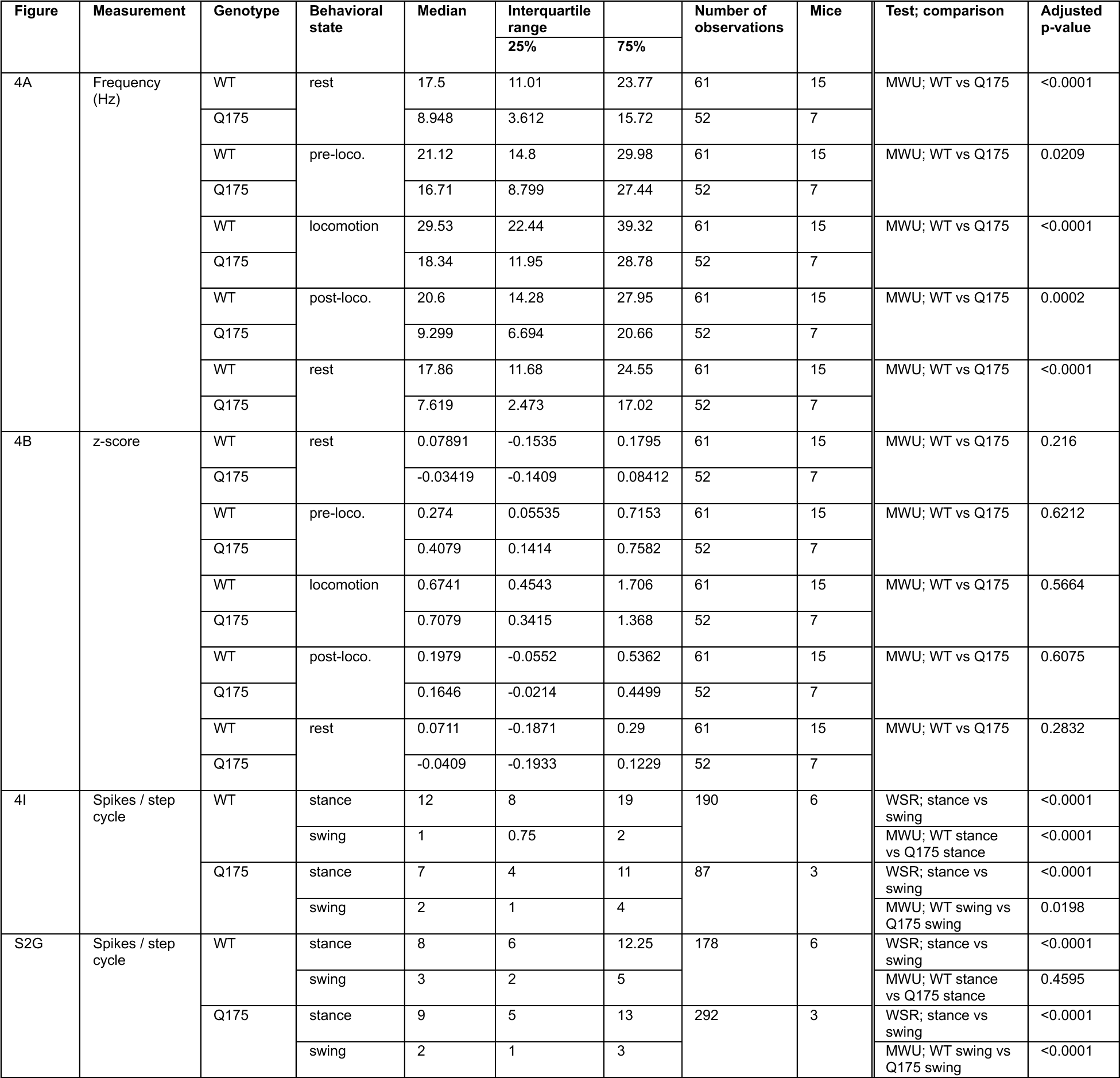
The frequencies of resting and locomotion-associated type 1 STN activity are reduced in Q175 mice. Related to Figures 4 and S2.

| Figure | Measurement | Genotype | Behavioral state | Median | Interquartile range |  | Number of observations | Mice | Test; comparison | Adjusted p-value |
| --- | --- | --- | --- | --- | --- | --- | --- | --- | --- | --- |
|  |  |  |  |  | 25% | 75% |  |  |  |  |
| 4A | Frequency (Hz) | WT | rest | 17.5 | 11.01 | 23.77 | 61 | 15 | MWU; WT vs Q175 | <0.0001 |
|  |  | Q175 |  | 8.948 | 3.612 | 15.72 | 52 | 7 |  |  |
|  |  | WT | pre-loco. | 21.12 | 14.8 | 29.98 | 61 | 15 | MWU; WT vs Q175 | 0.0209 |
|  |  | Q175 |  | 16.71 | 8.799 | 27.44 | 52 | 7 |  |  |
|  |  | WT | locomotion | 29.53 | 22.44 | 39.32 | 61 | 15 | MWU; WT vs Q175 | <0.0001 |
|  |  | Q175 |  | 18.34 | 11.95 | 28.78 | 52 | 7 |  |  |
|  |  | WT | post-loco. | 20.6 | 14.28 | 27.95 | 61 | 15 | MWU; WT vs Q175 | 0.0002 |
|  |  | Q175 |  | 9.299 | 6.694 | 20.66 | 52 | 7 |  |  |
|  |  | WT | rest | 17.86 | 11.68 | 24.55 | 61 | 15 | MWU; WT vs Q175 | <0.0001 |
|  |  | Q175 |  | 7.619 | 2.473 | 17.02 | 52 | 7 |  |  |
| 4B | z-score | WT | rest | 0.07891 | -0.1535 | 0.1795 | 61 | 15 | MWU; WT vs Q175 | 0.216 |
|  |  | Q175 |  | -0.03419 | -0.1409 | 0.08412 | 52 | 7 |  |  |
|  |  | WT | pre-loco. | 0.274 | 0.05535 | 0.7153 | 61 | 15 | MWU; WT vs Q175 | 0.6212 |
|  |  | Q175 |  | 0.4079 | 0.1414 | 0.7582 | 52 | 7 |  |  |
|  |  | WT | locomotion | 0.6741 | 0.4543 | 1.706 | 61 | 15 | MWU; WT vs Q175 | 0.5664 |
|  |  | Q175 |  | 0.7079 | 0.3415 | 1.368 | 52 | 7 |  |  |
|  |  | WT | post-loco. | 0.1979 | -0.0552 | 0.5362 | 61 | 15 | MWU; WT vs Q175 | 0.6075 |
|  |  | Q175 |  | 0.1646 | -0.0214 | 0.4499 | 52 | 7 |  |  |
|  |  | WT | rest | 0.0711 | -0.1871 | 0.29 | 61 | 15 | MWU; WT vs Q175 | 0.2832 |
|  |  | Q175 |  | -0.0409 | -0.1933 | 0.1229 | 52 | 7 |  |  |
| 4I | Spikes / step cycle | WT | stance | 12 | 8 | 19 | 190 | 6 | WSR; stance vs swing | <0.0001 |
|  |  |  | swing | 1 | 0.75 | 2 |  |  | MWU; WT stance vs Q175 stance | <0.0001 |
|  |  | Q175 | stance | 7 | 4 | 11 | 87 | 3 | WSR; stance vs swing | <0.0001 |
|  |  |  | swing | 2 | 1 | 4 |  |  | MWU; WT swing vs Q175 swing | 0.0198 |
| S2G | Spikes / step cycle | WT | stance | 8 | 6 | 12.25 | 178 | 6 | WSR; stance vs swing | <0.0001 |
|  |  |  | swing | 3 | 2 | 5 |  |  | MWU; WT stance vs Q175 stance | 0.4595 |
|  |  | Q175 | stance | 9 | 5 | 13 | 292 | 3 | WSR; stance vs swing | <0.0001 |
|  |  |  | swing | 2 | 1 | 3 |  |  | MWU; WT swing vs Q175 swing | <0.0001 |

**Table S5.**
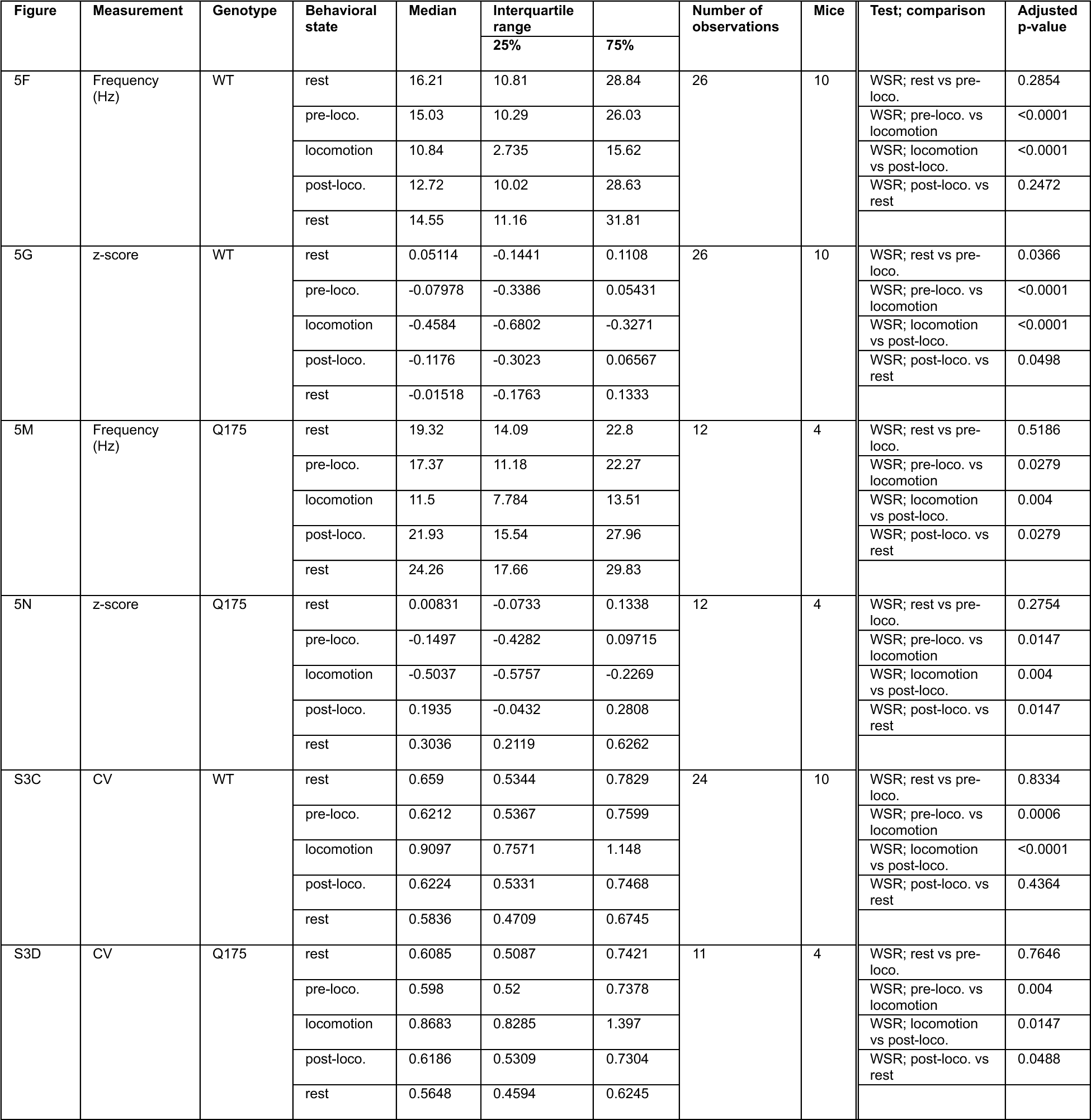
Type 2 STN neurons in WT and Q175 mice exhibit locomotion-associated decreases in firing. Related to Figures 5 and S3.

**Table S6.** The frequencies and patterns of resting and locomotion-associated type 2 STN neuron activity are similar in WT and Q175 HD mice. Related to Figure 6.

| Figure | Measurement | Genotype | Behavioral state | Median | Interquartile range |  | Number of observations | Mice | Test; comparison | Adjusted p-value |
| --- | --- | --- | --- | --- | --- | --- | --- | --- | --- | --- |
|  |  |  |  |  | 25% | 75% |  |  |  |  |
| 6A | Frequency (Hz) | WT | rest | 16.21 | 10.81 | 28.84 | 26 | 10 | MWU; WT vs Q175 | 0.8648 |
|  |  | Q175 |  | 19.32 | 14.09 | 22.8 | 12 | 4 |  |  |
|  |  | WT | pre-loco. | 15.03 | 10.29 | 26.03 | 26 | 10 | MWU; WT vs Q175 | 0.9877 |
|  |  | Q175 |  | 17.37 | 11.18 | 22.27 | 12 | 4 |  |  |
|  |  | WT | locomotion | 10.84 | 2.735 | 15.62 | 26 | 10 | MWU; WT vs Q175 | 0.5663 |
|  |  | Q175 |  | 11.5 | 7.784 | 13.51 | 12 | 4 |  |  |
|  |  | WT | post-loco. | 12.72 | 10.02 | 28.63 | 26 | 10 | MWU; WT vs Q175 | 0.5663 |
|  |  | Q175 |  | 21.93 | 15.54 | 27.96 | 12 | 4 |  |  |
|  |  | WT | rest | 14.55 | 11.16 | 31.81 | 26 | 10 | MWU; WT vs Q175 | 0.2328 |
|  |  | Q175 |  | 24.26 | 17.66 | 29.83 | 12 | 4 |  |  |
| 6B | z-score | WT | rest | 0.05114 | -0.1441 | 0.1108 | 26 | 10 | MWU; WT vs Q175 | 0.9383 |
|  |  | Q175 |  | 0.00831 | -0.0733 | 0.1338 | 12 | 4 |  |  |
|  |  | WT | pre-loco. | -0.07978 | -0.3386 | 0.05431 | 26 | 10 | MWU; WT vs Q175 | 0.9877 |
|  |  | Q175 |  | -0.1497 | -0.4282 | 0.09715 | 12 | 4 |  |  |
|  |  | WT | locomotion | -0.4584 | -0.6802 | -0.3271 | 26 | 10 | MWU; WT vs Q175 | 0.6311 |
|  |  | Q175 |  | -0.5037 | -0.5757 | -0.2269 | 12 | 4 |  |  |
|  |  | WT | post-loco. | -0.1176 | -0.3023 | 0.06567 | 26 | 10 | MWU; WT vs Q175 | 0.0178 |
|  |  | Q175 |  | 0.1935 | -0.0432 | 0.2808 | 12 | 4 |  |  |
|  |  | WT | rest | -0.01518 | -0.1763 | 0.1333 | 26 | 10 | MWU; WT vs Q175 | 0.0008 |
|  |  | Q175 |  | 0.3036 | 0.2119 | 0.6262 | 12 | 4 |  |  |
| 6I | Spikes / step cycle | WT | stance | 4 | 2 | 7 | 180 | 3 | WSR; stance vs swing | <0.0001 |
|  |  |  | swing | 1 | 0 | 2 |  |  | MWU; WT stance vs Q175 stance | 0.1597 |
|  |  | Q175 | stance | 6 | 3 | 7 | 24 | 2 | WSR; stance vs swing | <0.0001 |
|  |  |  | swing | 2 | 1 | 2.75 |  |  | MWU; WT swing vs Q175 swing | 0.0602 |

**Table S7.**
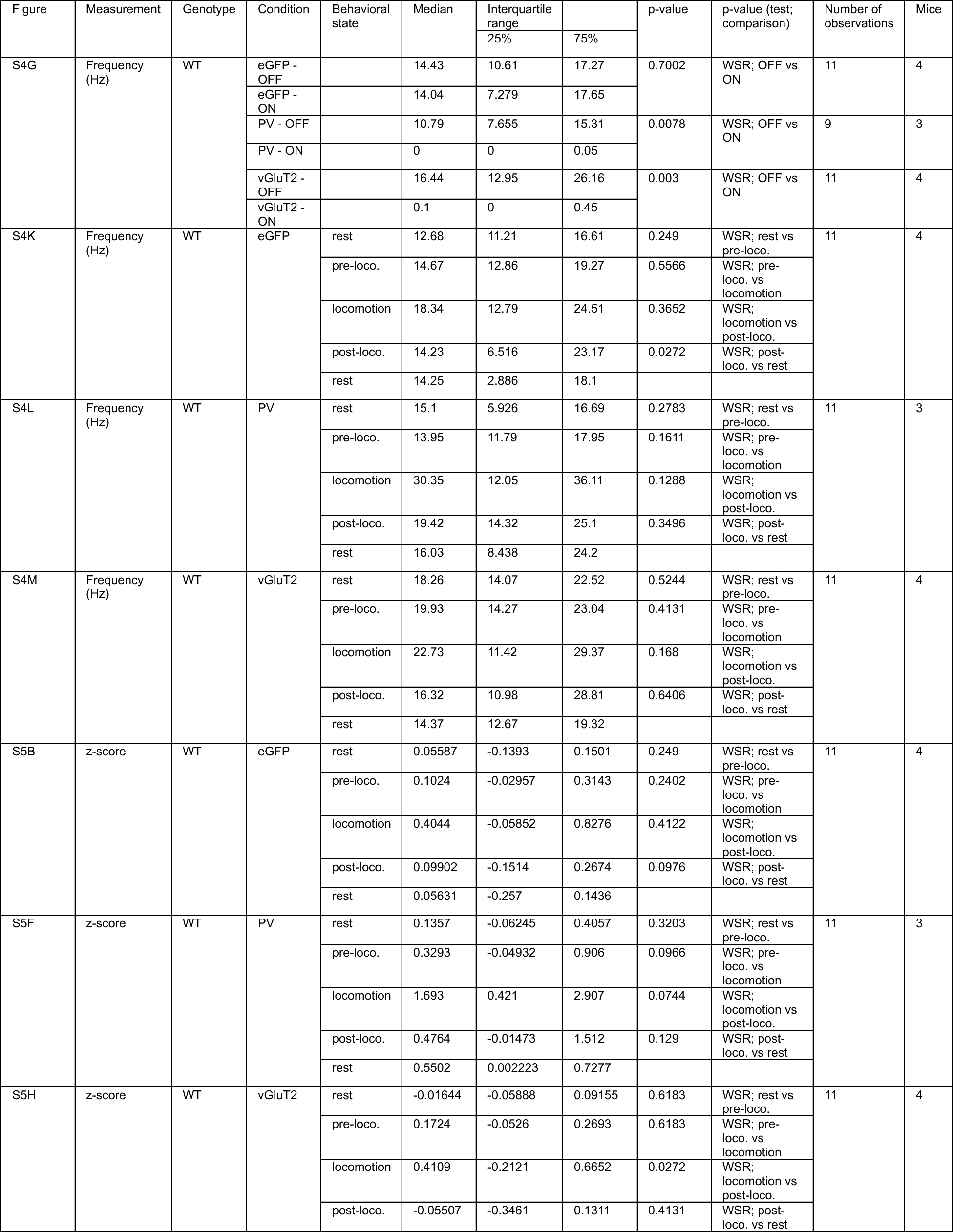
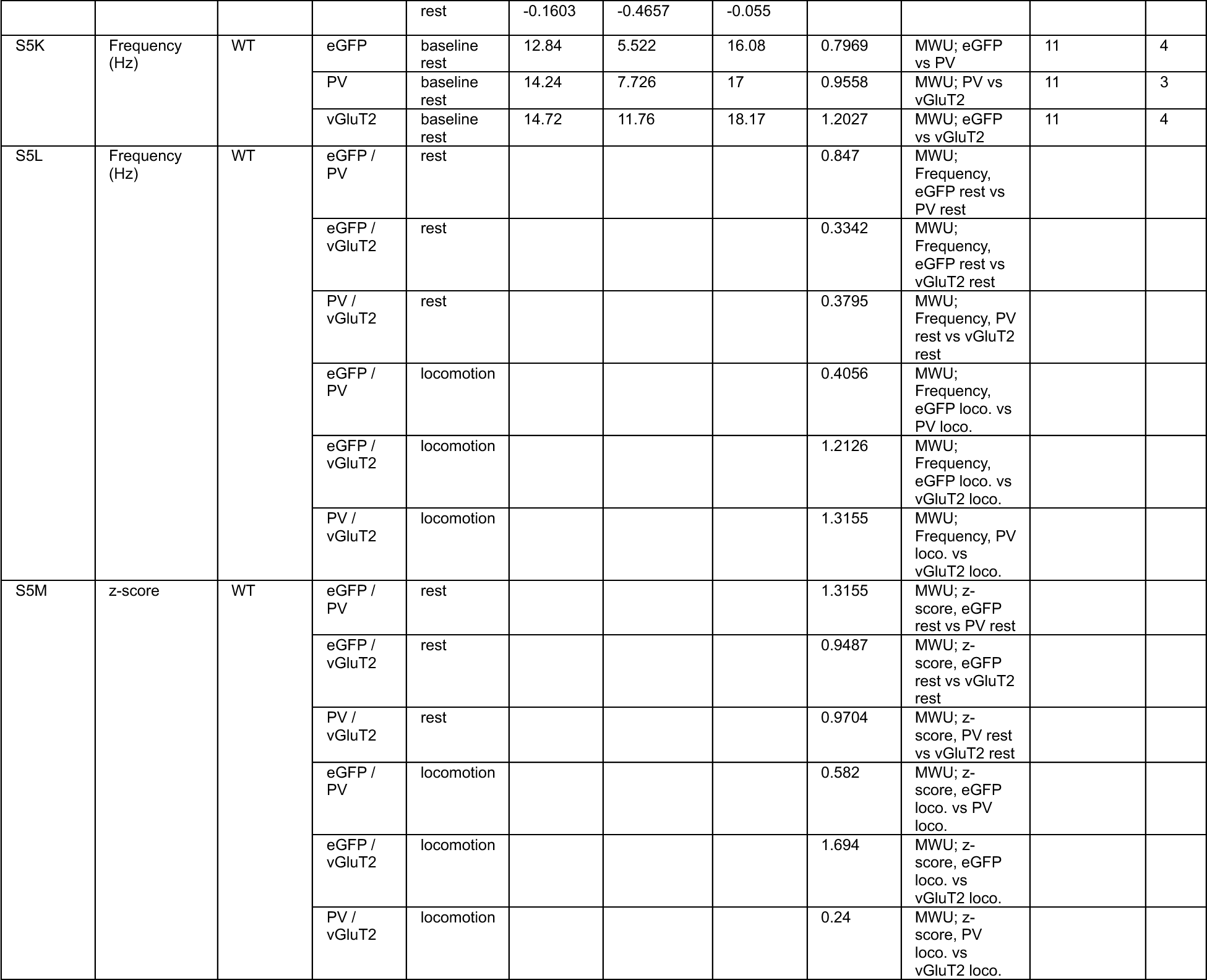
PV+ and vGluT2+ STN neurons exhibit heterogeneous locomotion encoding properties. Related to Figures S4 and S5.

**Table S8.** Optogenetic inhibition of movement-related STN activity dysregulates locomotion. Related to Figures 7 and S6.

| Figure | Measurement | Genotype | Condition | Median | Interquartile range |  | Number of observations | Mice | Test; comparison | Adjusted p-value |
| --- | --- | --- | --- | --- | --- | --- | --- | --- | --- | --- |
|  |  |  |  |  | 25% | 75% |  |  |  |  |
| 7B | Frequency (Hz) | WT | NpHR - Medial - OFF | 13.75 | 10.82 | 19.18 | 24 | 4 | WSR; Medial, OFF vs ON | <0.0001 |
|  |  |  | NpHR - Medial - ON | 0.05 | 0 | 0.15 |  |  | WSR; Lateral, OFF vs ON | <0.0001 |
|  |  |  | NpHR - Lateral - OFF | 19.75 | 14.51 | 24.68 | 34 | 5 | MWU; Med. OFF vs Lat. OFF | 0.0608 |
|  |  |  | NpHR - Lateral - ON | 0.1 | 0 | 1.825 |  |  | MWU; Med. ON vs Lat. ON | 0.0921 |
| S6E | Frequency (Hz) | WT | NpHR - CaMKII - OFF | 19.36 | 12.73 | 24.22 | 36 | 4 | WSR; CaMKII, OFF vs ON | <0.0001 |
|  |  |  | NpHR - CaMKII - ON | 0.05 | 0 | 0.7625 |  |  | WSR; vGluT2, OFF vs ON | <0.0001 |
|  |  |  | NpHR - vGluT2 - OFF | 15.03 | 11.35 | 18.7 | 22 | 3 | MWU; CaMKII OFF vs vGluT2 OFF | 0.1324 |
|  |  |  | NpHR - vGluT2 - ON | 0.075 | 0 | 0.325 |  |  | MWU; CaMKII ON vs vGluT2 ON | 0.7504 |
| S6F | Frequency (Hz) | WT | Type 1 - NpHR - OFF | 18.19 | 13.74 | 26.24 | 13 | 4 | WSR; Type 1, OFF vs ON | 0.0002 |
|  |  |  | Type 1 - NpHR - ON | 0.1 | 0 | 0.425 |  |  |  |  |
|  |  |  | Type 2 - NpHR - OFF | 19.64 | 12.12 | 24.48 | 6 | 3 | WSR; Type 2, OFF vs ON | 0.0312 |
|  |  |  | Type 2 - NpHR - ON | 0 | 0 | 0.05 |  |  |  |  |
|  |  |  | uncorr. - NpHR - OFF | 6.635 | 6.381 | 7.935 | 3 | 2 | WSR; uncorr., OFF vs ON | n/a |
|  |  |  | uncorr. - NpHR - ON | 0 | 0 | 0 |  |  |  |  |
| 7D | $\Delta$ velocity (cm/s) | WT | eGFP | 0.000061 | -0.0015 | 0.00159 | 121 | 4 | MWU; eGFP vs NpHR Med. | 1.1949 |
|  |  |  | NpHR - Medial | -0.000216 | -0.00278 | 0.00154 | 42 | 4 | MWU; NpHR Med. vs NpHR Lat. | 1.2136 |
|  |  |  | NpHR - Lateral | -0.000061 | -0.00226 | 0.00153 | 115 | 6 | MWU; eGFP vs NpHR Lat. | 0.6643 |
| 7K | Locomotor bout duration (s) | WT | eGFP | 3.537 | 2.014 | 6.102 | 40 | 4 | MWU; eGFP vs NpHR Med. | 0.4021 |
|  |  |  | NpHR - Medial | 2.643 | 1.158 | 7.806 | 25 | 4 | MWU; NpHR Med. vs NpHR Lat. | 0.2016 |
|  |  |  | NpHR - Lateral | 1.839 | 0.727 | 3.058 | 41 | 6 | MWU; eGFP vs NpHR Lat. | 0.0006 |
| S6G | Locomotor bout duration (s) | WT | eGFP | 3.537 | 2.014 | 6.102 | 40 | 4 | MWU; eGFP vs NpHR Med. | 0.4021 |
|  |  |  | NpHR - Medial | 2.643 | 1.158 | 7.806 | 25 | 4 | MWU; eGFP vs NpHR Cen. | 0.0048 |
|  |  |  | NpHR - Central | 1.839 | 1.136 | 2.403 | 17 | 4 | MWU; eGFP vs NpHR Lat. | 0.005 |
|  |  |  | NpHR - Lateral | 1.88 | 0.2725 | 3.336 | 24 | 5 |  |  |
| S6H | Locomotor bout duration (s) | WT | NpHR - CaMKII | 1.684 | 0.352 | 2.492 | 29 | 4 | MWU; CaMKII vs vGluT2 | 0.0676 |
|  |  |  | NpHR - vGluT2 | 2.057 | 1.367 | 4.271 | 12 | 3 |  |  |
| S6I | Locomotor bout duration (s) | WT | NpHR - Early trials | 1.963 | 0.69 | 3.655 | 13 | 6 | WSR; NpHR, Early vs Late | 0.8926 |
|  |  |  | NpHR - Late trials | 2.028 | 0.468 | 5.399 | 13 | 6 |  |  |
| 7L | $\Delta$ Stance time (s) | WT | eGFP | 0 | -0.055 | 0.0981 | 21 | 4 | MWU; eGFP vs NpHR | 0.0085 |
|  |  |  | NpHR | 0.135 | 0.07813 | 0.175 | 17 | 4 |  |  |
| 7M | $\Delta$ Stance speed (cm/s) | WT | eGFP | -0.4987 | -1.424 | 0.3295 | 21 | 4 | MWU; eGFP vs NpHR | 0.0019 |
|  |  |  | NpHR | -3.432 | -5.27 | -1.459 | 17 | 4 |  |  |
| 7N | $\Delta$ Time between strides (s) | WT | eGFP | 0.0625 | -0.08 | 0.1215 | 18 | 4 | MWU; eGFP vs NpHR | 0.0132 |

